# Cytokinin inhibits fungal development and virulence by targeting the cytoskeleton and cellular trafficking

**DOI:** 10.1101/2020.11.04.369215

**Authors:** Rupali Gupta, Gautam Anand, Lorena Pizarro, Dana Laor, Neta Kovetz, Noa Sela, Tal Yehuda, Ehud Gazit, Maya Bar

**Affiliations:** Department of Plant Pathology and Weed Research, ARO, Volcani Center, Rishon LeZion 7505101, Israel; Institute of Agri-food, Animal and Environmental Sciences, Universidad de O’Higgins, Chile; The Shmunis School of Biomedicine and Cancer Research, Tel Aviv University, Tel Aviv 6997801, Israel

**Keywords:** Cytokinin, cell cycle, cytoskeleton, endocytosis, *Botrytis cinerea*, *Saccharomyces cerevisiae*

## Abstract

Cytokinin (CK) is an important plant developmental regulator, having activities in many aspects of plant life and its response to the environment. CKs are involved in diverse processes in the plant, including stem-cell maintenance, vascular differentiation, growth and branching of roots and shoots, leaf senescence, nutrient balance and stress tolerance. In some cases, phytopathogens secrete CKs. It has been suggested that to achieve pathogenesis in the host, CK-secreting biotrophs manipulate CK signaling to regulate the host cell cycle and nutrient allocation. CK is known to induce host plant resistance to several classes of phytopathogens from a handful of works, with induced host immunity *via* salicylic acid signaling suggested to be the prevalent mechanism for this host resistance.

Here, we show that CK directly inhibits the growth, development, and virulence of fungal phytopathogens. Focusing on *Botrytis cinerea (Bc)*, we demonstrate that various aspects of fungal development can be reversibly inhibited by CK. We also found that CK affects both budding and fission yeast in a similar manner. Investigating the mechanism by which CK influences fungal development, we conducted RNA-NGS on mock and CK treated *B. cinerea* samples, finding that CK inhibits the cell cycle, cytoskeleton, and endocytosis. Cell biology experiments demonstrated that CK affects cytoskeleton structure and cellular trafficking in *Bc*, lowering endocytic rates and endomembrane compartment sizes, likely leading to reduced growth rates and arrested developmental programs. Mutant analyses in yeast confirmed that the endocytic pathway is altered by CK.

Our work uncovers a remarkably conserved role for a plant growth hormone in fungal biology, suggesting that pathogen-host interactions resulted in fascinating molecular adaptations on fundamental processes in eukaryotic biology.

**Importance:** Cytokinins (CKs), important plant growth/ developmental hormones, have previously been associated with host disease resistance. Here, we demonstrate that CK directly inhibits the growth, development, and virulence of *B. cinerea* (*Bc*) and many additional phytopathogenic fungi. Molecular and cellular analyses revealed that CK is not toxic to *Bc*, but rather, *Bc* likely recognizes CK and responds to it, resulting in cell cycle and individual cell growth retardation, via downregulation of cytoskeletal components and endocytic trafficking. Mutant analyses in yeast confirmed that the endocytic pathway is a CK target. Our work demonstrates a conserved role for CK in yeast and fungal biology, suggesting that suggesting that pathogen-host interactions may cause molecular adaptations on fundamental processes in eukaryotic biology.

## Introduction

Cytokinins (CKs) are a class of extensively studied plant hormones, well known for their involvement in various aspects of plant life (Mok & Mok, 2001; Sakakibara, 2006; Werner & Schmülling, 2009; Keshishian & Rashotte, 2015). Some findings have suggested a role for CKs in fungal pathogenesis (Walters & McRoberts, 2006; Babosha, 2009; Sharma *et al*., 2010; Choi *et al*., 2011). In some cases, plant pathogens can secrete CKs, or induce CK production in the host plant, possibly in order to achieve pathogenesis in the host (Jameson, 2000). Conidia, mycelia of some fungi, and germinating uredospores of *Puccinia graminis* and *P. recondite* have been shown to accumulate CK, manipulating CK signaling to regulate host plant cell cycle (Greene, 1980; Nieto & Frankenberger, 1991).

CK has also been shown to promote resistance to plant pathogens that do not secrete CK. High levels of CKs were found to increase the plants’ resistance to bacterial and fungal pathogens (Swartzberg *et al*., 2008; Choi *et al*., 2010; Grosskinsky *et al*., 2011; Ballaré, 2011; Gupta *et al*., 2020a,b). Different mechanisms have been suggested for this enhanced resistance. In Arabidopsis, it was suggested that CK-mediated resistance functions through salicylic acid (SA) dependent mechanisms (Choi *et al*., 2010). An additional study suggested that CK signaling enhances the contribution of SA-mediated immunity in hormone disease networks (Naseem *et al*., 2012). However, in tobacco, an SA-Independent, phytoalexin-dependent mechanism was suggested (Grosskinsky *et al*., 2011). We recently reported that CK induces systemic immunity in tomato (*Solanum lycopersicum*), promoting resistance to fungal and bacterial pathogens (Gupta *et al*., 2020a,b) including *Botrytis cinerea,* in an SA and ethylene dependent mechanism.

*B. cinerea* (grey mold), *Sclerotium rolfsii* (collar rot), and *Fusarium oxysporum* f. sp. *lycopersici* (fusarium wilt) are widespread fungal plant pathogens that infect hundreds of plant species and cause huge losses every year (Tsahouridou & Thanassoulopoulos, 2002; Williamson *et al*., 2007; Dean *et al*., 2012).

Given that CK can induce plant immunity and restrict phytopathogen growth in certain cases, direct effects of CK against phytopathogens are an intriguing possibility. A direct effect of CK on bacterial pathogens was ruled out in previous works (Naseem *et al*., 2012; Gupta *et al*., 2020a). Interestingly, CK did not affect germination and elongation of germ tubes, but strongly inhibited appressorium formation of *Erysiphe graminis*, an obligate biotrophic powdery mildew causing barley pathogen (Liu & Bushnell, 1986). Another work described an increase in germination of conidia of two *Erysiphe* powdery mildew pathogens, *E. graminis* and *E. cichoracearum*, in the presence of CK (Mishina *et al*., 2002). Fungal pathogens of the species *Erysiphe* are known to produce CKs. High levels of CK was also reported to inhibit mycelial growth and pathogenesis of fungi in canola (Sharma *et al*., 2010).

In this work, we investigate the direct effects of CK on fungal plant pathogens, demonstrating that CK directly inhibits the growth, development, and virulence of fungal plant pathogens. *B. cinerea* (*Bc*) growth, sporulation, and spore germination were all inhibited by CK, in the absence of a host plant. We found similar effects in a variety of plant pathogenic fungi. Molecular and cellular analyses revealed that CK is not toxic to *Bc*, but rather, *Bc* likely recognizes CK and responds to it, resulting in cell cycle and individual cell growth retardation. CK reduced *Bc* virulence in a reversible manner, confirming that no irreversible harm was caused to fungal development. RNAseq confirmed that CK downregulates the cell cycle and cytoskeleton, as well as cellular trafficking. Interestingly, we also found that CK affects two additional fungi from the ascomycota division that are not phytopathogens, but rather, yeasts. Both budding and fission yeast were inhibited by CK. Further to the effects observed in the RNAseq data, we confirmed that CK affects the cytoskeleton and cellular trafficking in *Bc*, mislocalizing actin filaments and lowering endocytic rates and endomembrane compartment sizes, likely underlying the reduced growth rates and arrested developmental programs. Mutant analyses in yeast confirmed that the endocytic pathway is affected by CK. Our work uncovers a novel, remarkably conserved role for a primary plant growth hormone in fungal biology, demonstrating that interaction between pathogen and host resulted in fascinating molecular adaptations on fundamental processes in eukaryotic biology. In time, this may hold promise for the development of CKs as antifungal agents in specific cases.

## Results

### Cytokinin inhibits disease caused by *B. cinerea* and *S. rolfsii*, but not *F. oxysporum* f. sp*. lycopersici*

We have previously reported that CK reduces tomato disease by inducing immunity (Gupta *et al*., 2020b). Fungal pathogens with different lifestyles have different infection and pathogenesis strategies, and host plants employ different protection mechanisms to resist different types of fungal pathogens. Three phytopathogenic fungi with varied lifestyles and infection modes were selected. *B. cinerea (Bc),* an airborne necrotrophic spore producing ascomycete, that causes gray mold disease in >1400 hosts (Fillinger & Elad, 2016); *S. rolfsii (Sr)*-a soilborne necrotrophic basidiomycete that does not produce spores, that causes southern blight disease in hundreds of hosts (Arnold, 2008); and *F. oxysporum* f. sp. *lycopersici (Fol)*, a soilborne necrotrophic ascomycete that is known to produce CK (Vrabka *et al*., 2018; Sørensen *et al*., 2018), and causes wilt disease in a host specific manner (Stravato *et al*., 1999). In order to examine the ability of CK to reduce disease caused by different fungal pathogens, we treated tomato plants with 100 µM CK (6-BAP) 24 hours prior to pathogen inoculation. **Fig. 1** details the effect of CK in tomato disease caused by different phytopathogens. 6-BAP pre-treatment significantly decreased disease levels caused by the necrotrophic fungal pathogens *Bc*, as we previously reported (Gupta *et al*., 2020b) (**Fig. 1a**), and *Sr* (**Fig. 1b**). Upwards of 50% disease reduction in tomato plants was observed with *Bc* and *Sr* as compared to control (**Fig. 1 a,b**). However, no disease reduction was observed with *Fol* (**Fig. 1c**).

**Fig. 1.**
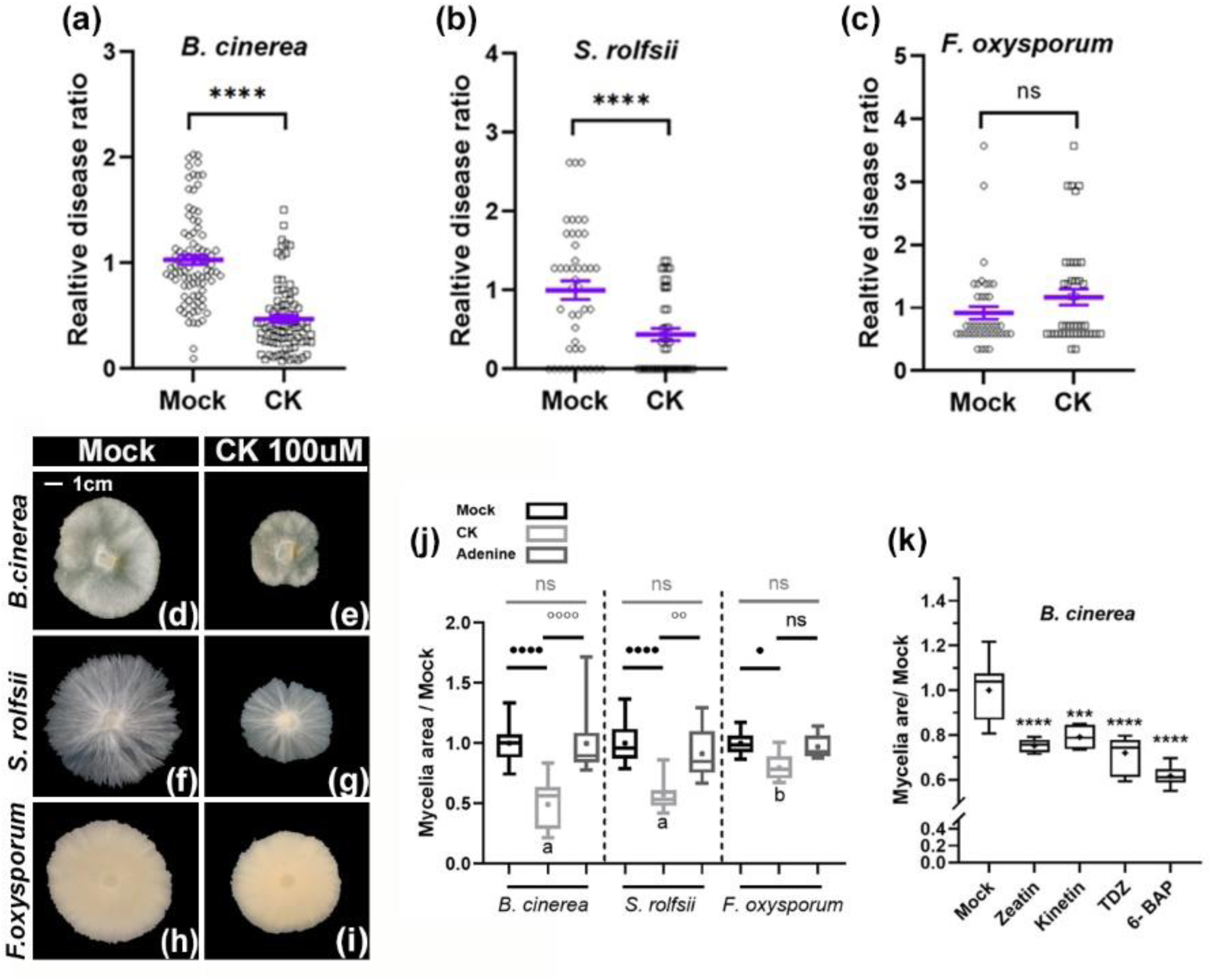
Direct effect of cytokinin on fungal growth and disease. **(a)-(c)** *S. lycopersicum* cv M82 leaves were treated with 100 µM CK (6-BAP; 6-Benzylaminopurine) and inoculated with indicated pathogens 24 hours later. Plant disease was quantified as indicated in the material and methods section, disease in the mock treatment was set to 1. Graphs represent results from 4-6 biological repeats ±SE, N>40. Asterisks represent significant difference among means in a two-tailed t-test, ****p<0.0001, ns-non-significant. All individual values plotted, purple bar represents mean ±SE. **(d)-(i)** *B. cinerea, S. rolfsii and F. oxysporum* were cultured on potato dextrose agar (PDA) plates in the presence of 100 µM 6-BAP. **(j)** Quantification of results from 4-6 biological repeats ±SE, including adenine (100 µM) as a control, N>20. Asterisks (differences between Mock and CK) and letters (differences between the level of CK growth inhibition in the different fungi) indicate significance in one-way ANOVA with a Bonferroni post hoc test, *p<0.05, **p<0.01, ****p<0.0001, ns=non-significant. **(k)** *B. cinerea* was cultured on potato dextrose agar (PDA) plates in the presence of 100 µM of the indicated CK compound (TDZ=Thidiazuron). Quantification of results from 3 biological repeats ±SE, N=12. Asterisks indicate significance in one-way ANOVA with a Tukey post hoc test, ***p<0.001, ****p<0.0001. **(i-k)** Dose response of *B. cinerea, S. rolfsii and F. oxysporum* to CK-different concentrations of 6-BAP as indicated. Graphs represent 3 biological repeats ±SE, N>6. Letters indicate significance in a one-way ANOVA, ****p<0.0001 in all cases, with a Tukey post-hoc test. **j-k**: Box-plot displays minimum to maximum values, with inner quartile ranges indicated by box and outer-quartile ranges by whiskers. Line indicates median, dot indicates mean.

Cytokinin directly inhibits *B. cinerea, S. rolfsii*, and *F. oxysporum* f. sp. *lycopersici*

In order to examine a possible direct effect of CK on fungal tomato pathogens, we used the above three phytopathogenic fungi. The effects of different CK concentrations and derivatives on the growth of *Bc, Sr* and *Fol* mycelia are shown in **Fig. 1**. The cyclic synthetic CK 6-BAP (**Fig. 1d-k**), the natural cyclic CKs zeatin and kinetin (**Fig. 1k**), and the synthetic bacterial derived non-cyclic CK thidiazurn (TDZ, **Fig. 1k**)- all inhibited the growth of *B. cinerea*. 6-BAP (6-benzyl amino purine) inhibited the growth of *Bc* in a dose-dependent manner. Approximately 50% inhibition was observed at 100 µM (**Fig. S1a**). *Sr* did not show growth reduction at 1 µM but growth inhibition at 10 µM and 100 µM was similar to *Bc* (**Fig. S1b**). The least effect of 6-BAP was observed on *Fol* (**Fig. S1c**), which was not inhibited by CK treatment *in planta* (**Fig. 1c**).

To examine the breadth of this phenomenon, we tested *in vitro* growth of additional phytopathogenic fungi in the presence of 100 µM 6-BAP or the control Adenine. Our results show that CK directly inhibits mycelial growth of fungal pathogens from several different classes (ascomycetes, basidiomycetes) and different lifestyles (hemibiotrophs, necrotrophs) (**Fig. S2**). All classes of phytopathogenic fungi tested were inhibited by CK (**Fig. S2a**), however, the level of inhibition also differed significantly among them, with *Fusarium* spp. showing the least inhibition (**Fig. S2b**). The phylogeny is detailed in **Fig. S2c**, and does not indicate that the ability to be inhibited by CK is specific to any particular class or taxon.

### Cytokinin inhibits *B. cinerea* sporulation, spore germination, and germ tube elongation

We studied the effect of CK fungal development: sporulation, spore germination, and germ tube elongation. In samples treated with 100 nM CK, there were approximately 50% less spores produced as compared to mock (**Fig. 2a,b**). The effect of CK on spore germination was even more pronounced (**Fig. 2c,d**). Less than 50% of the spores germinated with 100 nM CK. Interestingly, though less spores germinated in the presence of 6-BAP, the spores that did germinate had accelerated germ tube growth in 100 nM 6-BAP, and inhibited germ tube growth 100 µM 6-BAP (**Fig. 2c,e**). After 8 hours of growth with 100 µM 6-BAP, the germ tube length was 50% of the control (**Fig. 2e**), correlating with ∼50% inhibition in mycelial growth (**Fig. 1d,e,j,k**).

**Fig. 2.**
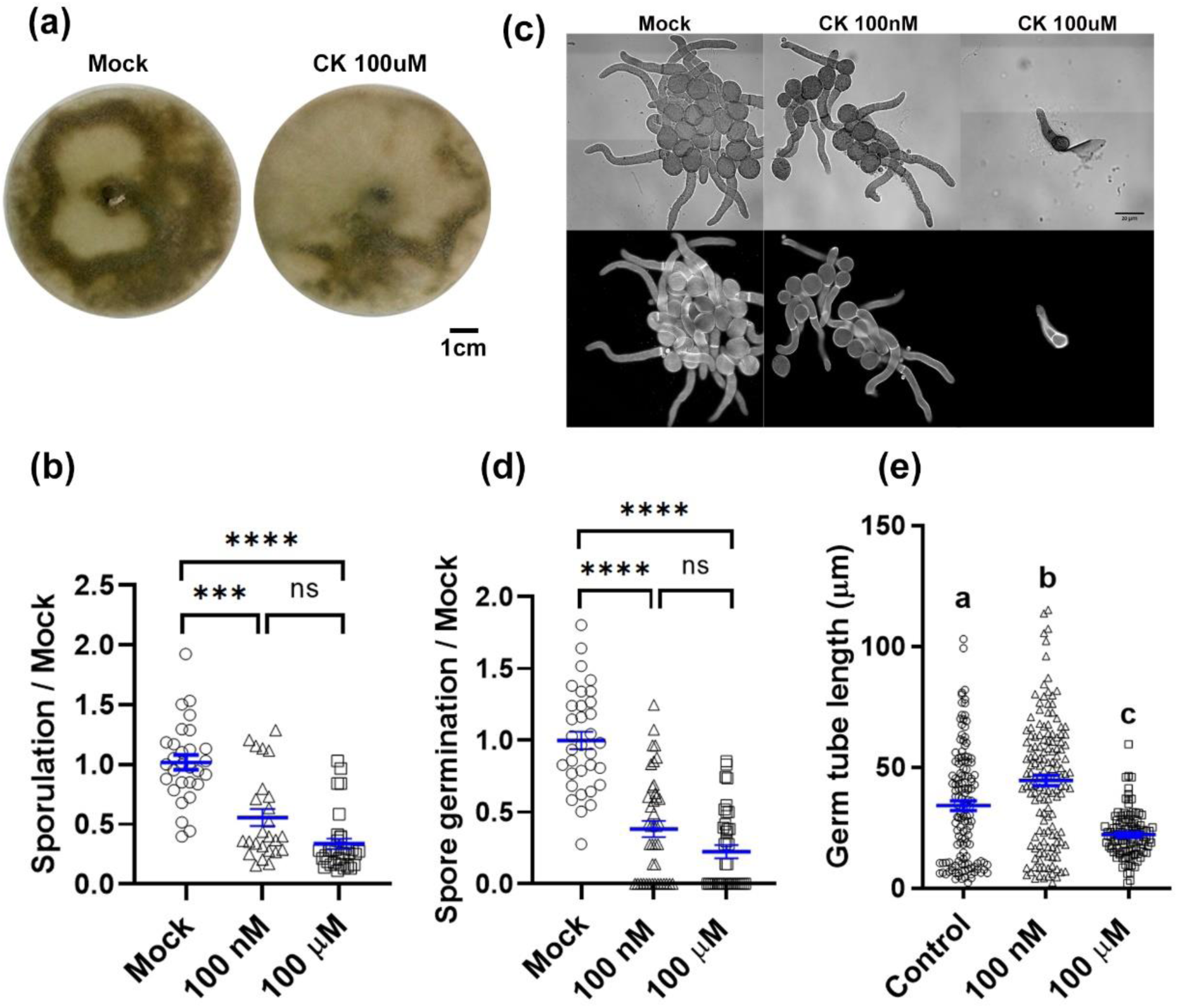
Cytokinin inhibits *Bc* sporulation and spore germination. **(a)-(b)** *B. cinerea* was cultured on PDA plates or PDB liquid broth in the presence of 100 nM or 100 µM 6-Benzylaminopurine (6-BAP). Spore formation is indicated by dark color **(a)** and quantified in **(b)**. Spore germination and germ tube elongation are demonstrated in **(c)** with calcofluor staining (scale bar = 20 µM), and quantified in **(d)-(e)**. **(b,d,e)** Quantification of results from 4 biological repeats ±SE, N>25. Asterisks or letters indicate significance in one-way ANOVA with a Tukey post hoc test, ***p<0.005, ****p<0.001. Individual values graphed, blue bar represents mean ±SE.

LC/MS Hormonal measurements in mature tomato leaves demonstrate that they can contain close to 100 ng/g active CKs (**Fig. S3**), as was previously reported for tomato leaves (Ghanem *et al*., 2008; Žižková *et al*., 2015). This amount roughly corresponds, depending on the CK derivative, to 400-500 nM, which did not affect mycelial growth of the phytopathogenic fungi we tested (**Fig. S1**), but did inhibit sporulation and spore germination in *B. cinerea* (**Fig. 2b-d**).

### Cytokinin reversibly attenuates *B. cinerea* virulence

Is direct CK inhibition of phytopathogenic fungi reversible? Or does CK irreversibly harm fungal development? Furthermore, since some aspects of *Bc* development are inhibited in CK concentrations normally found within plant leaves (**Figs. 2, S3**), how does CK affect fungal virulence? To answer these questions, we harvested *Bc* spores from fungi grown with or without CK, normalized the spore count to 10^5^ spores/mL, and used equal amounts of spores to infect tomato leaves. When spores harvested from mycelia grown with 6-BAP were used for infecting tomato leaves, no reduction in lesion size was observed as compared with spores grown without CK (**Fig. 3a**). We harvested spores from mycelia grown without CK, mixing half the spores with CK just prior to tomato leaf infection. When spores were treated with CK prior to infection, there was approximately 40% reduction in lesion area (**Fig. 3b**), comparable to disease reduction observed when treating plants with CK (**Fig. 1a**).

**Fig. 3.**
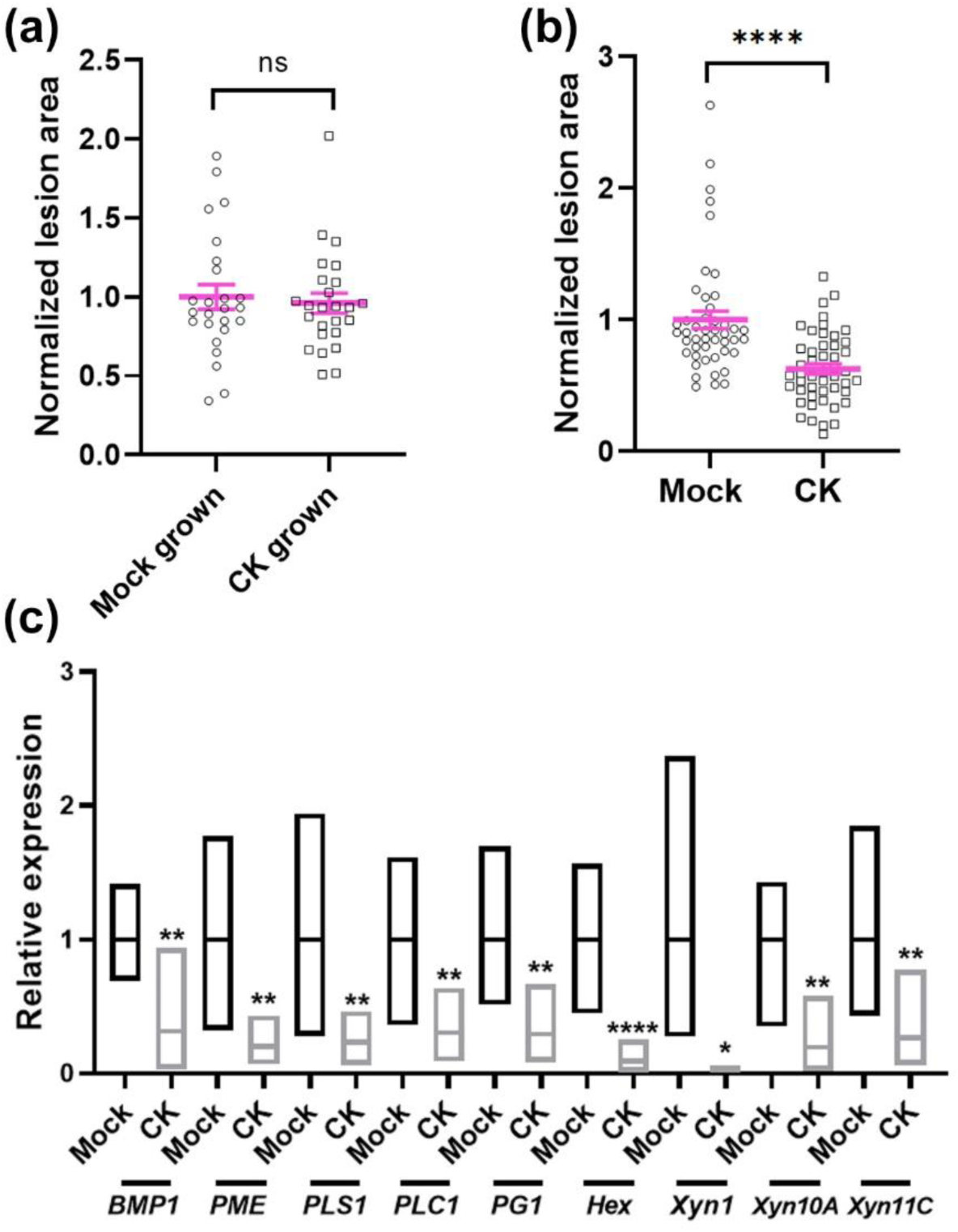
Cytokinin inhibits *B. cinerea* virulence. **(a)** Infectivity of spores sporulated from mycelia grown with or without cytokinin (CK; 6-Benzylaminopurine). Spores were harvested in a glucose/potassium media as described in the materials sections, and diluted to an equal concentration (the CK grown spores were concentrated as CK grown fungi produce reduced numbers). **(b)** Infectivity of spores grown in rich media and harvested from the same plate; half the spores were mixed with CK prior to infection. **(c)** *B. cinerea* was grown in PDB with the addition of 100 µM CK (6-BAP) or without (Mock). qRT-PCR was carried out on the virulence genes *BMP1, PME, PLS1, PLC1, PG1, hex, XynI, Xyn10A* and *Xyn11C*. Mock was set to 1. Gene expression values were normalized to a geometric mean of the expression of 3 housekeeping genes: Ubiquitin-conjugating enzyme E2, Iron-transport multicopper oxidase, and Adenosine deaminase. Graph represents 3 biological repeats, N=9. **a-b:** individual values graphed, pink bar represents mean ±SE. Results were analyzed for statistical significance using a two-tailed t-test, ****p<0.0001; ns-non-significant. **c:** floating bars, line indicates mean. Results were analyzed for statistical significance using Welch’s Anova with a Dunnett post hoc test, *p<0.05, **p<0.01, ****p<0.0001.

CK differentially down-regulated the expression levels of virulence genes in *Bc* (**Fig. 3c**). All tested genes related to virulence viz., *BcPG1* (endopolygalacturonase)*, BcXynl, BcXyn10A* and *BcXyn11C* (Xylanases), *BcPME1* (pectin methylesterase)*, BcPLC1* (phospholipase C), *BcBMP1* (mitogen-activated protein (MAP) kinase) and *BcPLS1* (tetraspanin) were expressed at significantly lower levels upon growth in the presence of 100 µM of 6-BAP. In addition, the expression of the woronin body protein *Bchex*, known to be involved in fungal membrane integrity (Torres-Ossandón *et al*., 2019), was down-regulated upon CK treatment. Gene expression was normalized to a geometric mean of 3 housekeeping genes: ubiquitin-conjugating enzyme E2 (*ubce*) (Silva-Moreno *et al*., 2016), Iron-transport multicopper oxidase, and Adenosine deaminase (Llanos *et al*., 2015).

### Cytokinin is not toxic to *B. cinerea*

Why is CK inhibiting fungal growth and development? Spores generated from fungus grown in the presence of CK were able to infect tomato leaves normally, once removed from the CK containing environment, indicating that the spores themselves were normal, and that disease reduction stems from a reduction in spore formation, spore germination, and germ tube growth (**Fig. 3**). To confirm that CK is not killing fungal cells, we examined fungal cellular leakage in the presence of CK. As shown in **Fig. 4a,b**, no significant leakage of nucleic acids and proteins were observed after 24 h treatment with 100 µM 6-BAP, when compared to the control. Additionally, no change in electrical conductivity was observed upon 24 hours of 100 µM 6-BAP treatment (**Fig. 4c)**. The high background measurements with CK alone are likely due to the presence of the aromatic ring structure. These results confirm that CK is not toxic to *Bc*.

**Fig. 4.**
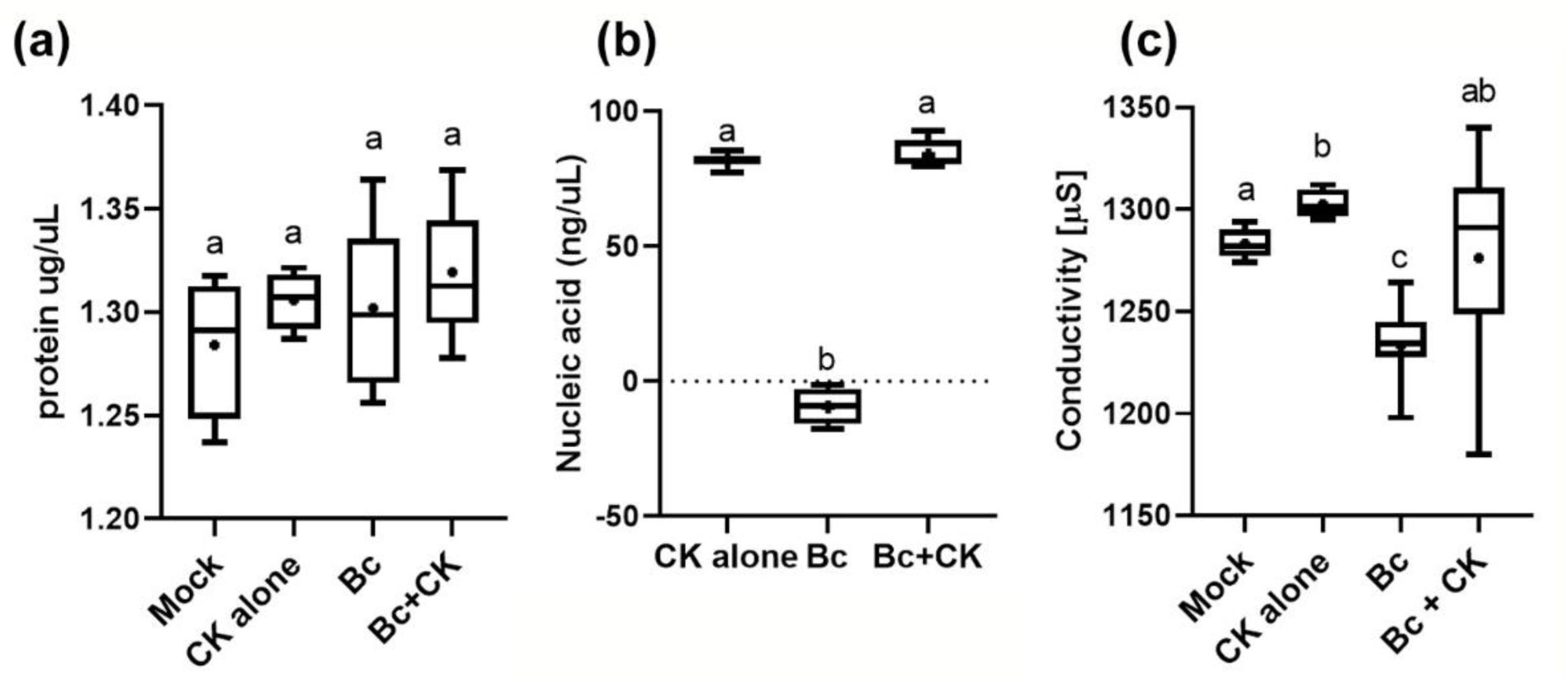
Cytokinin is not toxic to *B. cinerea*. *B. cinerea* (*Bc*) was cultured in PDB liquid broth, with or without 100 µM CK (6-Benzylaminopurine). After 24 hours, protein leakage **(a)**, nucleic acid leakage **(b)**, and media conductivity **(c)** were measured. Graphs represent 3 biological repeats ±SE, N>6. Letters indicate significance in one-way ANOVA with a Tukey (a) or Bonferroni (b,c) post hoc test; a p=ns; b,c p<0.0001. Box-plot with 2.5% whiskers; line indicates median, dot indicates mean. No significant difference between control media containing CK (without *Bc*) or *Bc* with CK was observed in any of the parameters.

### Transcriptome profiling reveals pathways affected by CK in *B. cinerea*

To gain insight into the effects CK has on the fungus, we conducted transcriptome profiling on *Bc* samples prepared from fungi grown with and without CK. Principle component analysis demonstrated that the biological replicates were clustered well together (**Fig. 5a**), with mock samples being very similar, and CK samples clustering together across PC1 (75%). The comparison yielded two clusters, exemplified in a heatmap (**Fig. 5b**): genes downregulated by CK when compared with mock (bottom cluster, marked in red) and genes upregulated by CK when compared with mock (top cluster, marked in green). Individual genes having a log_(2)_Fold-change of |2| or greater are provided in **Supplementary Data 1**. Interestingly, out of the 660 differentially expressed genes (DEGs), 470 were down regulated, indicating that CK has more of a suppressive effect on the *Bc* transcriptome. Distribution of DEGs into various biological pathways in Kyoto Encyclopedia of Genes and Genomes (KEGG) also supports this. Downregulated KEGG pathways included various pathways related to the cell cycle and DNA replication (**Fig. 5c**), as well as endocytosis, MAPK signaling, and a variety of metabolic pathways (**Fig. 5c**). The full KEGG list with adjusted p-values for each pathway is provided in **Supplementary data 2**. Upregulated KEGG pathways included various pathways related to protein biosynthesis and processing (**Fig. 5d**), as well as the peroxisome and phagosome (**Fig. 5d**). The full KEGG list with adjusted p-values for each pathway is provided in **Supplementary data 3**.

**Fig. 5.**
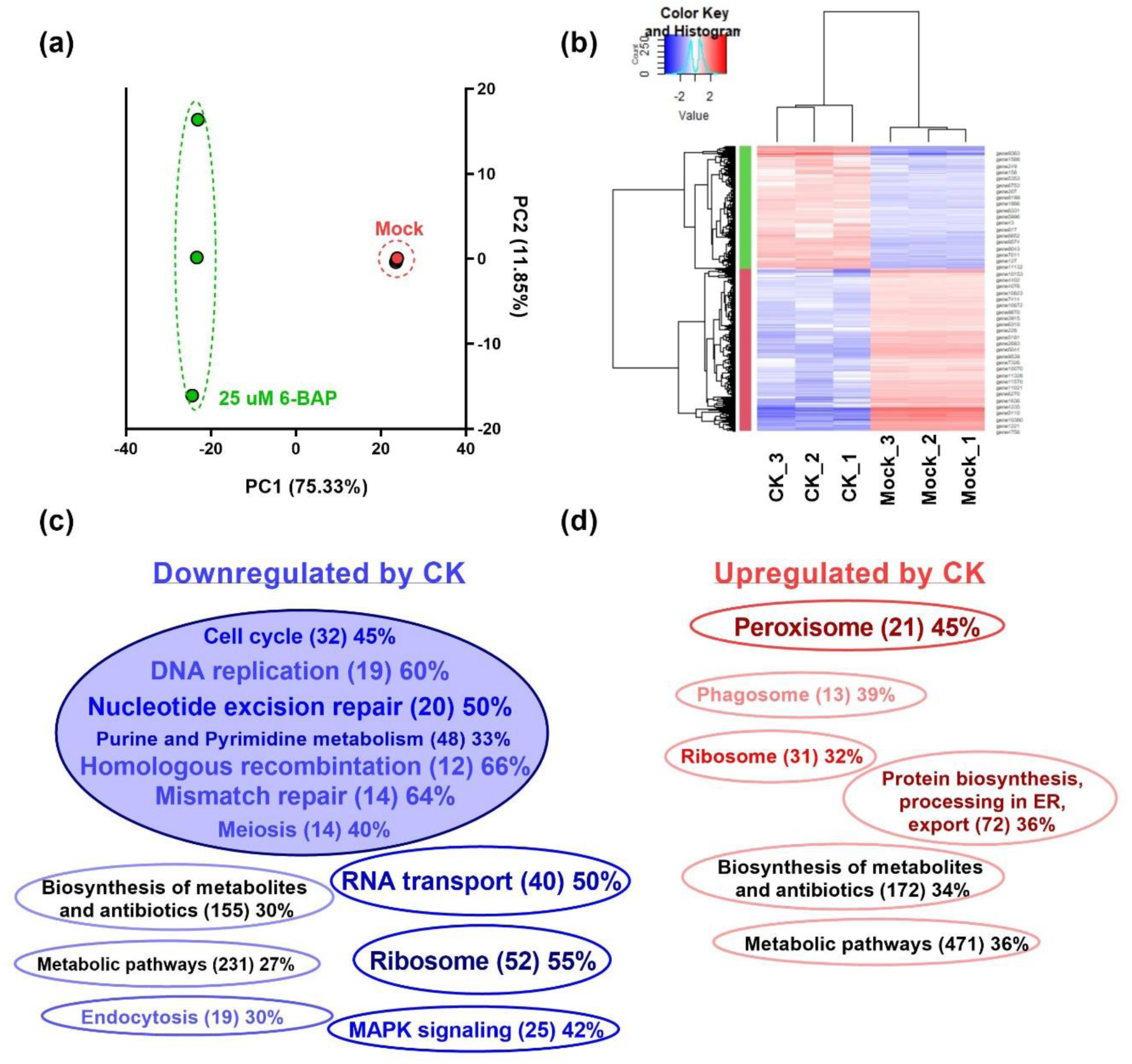
Transcriptomics reveal fungal pathways affected by CK. Analysis of Illumina Hiseq NGS of *Bc* samples Mock treated or CK treated (25 uM), 3 biological repeats each. Gene expression values were computed as FPKM. Differential expression analysis was completed using the DESeq2 R package. Genes with an adjusted p-value of no more than 0.05 and log2FC greater than 1 or lesser than -1 were considered differentially expressed. **(a)** Principle component analysis (PCA) of 3 biological repeats from each treatment. PCA was calculated using the R function prcomp. **(b)** Heatmap depicting the clustering of the different samples in terms of differentially expressed genes. Blue= negatively regulated by CK, red= positively regulated by CK. Color saturation indicates strength of differential expression. Heatmap visualization was performed using R Bioconductor. See also Supplemental data 1. **(c-d)** Analysis of statistically enriched pathways downregulated (c, blue) and upregulated (d, red) by CK. KOBAS 3.0 tool was used to detect the statistical enrichment of differential expression genes in Kyoto Encyclopedia of Genes and Genomes (KEGG) pathways and Gene Ontology (GO). Pathways were tested for significant enrichment using Fisher’s exact test, with Benjamini and Hochberg FDR correction. Corrected p-value was deemed significant at p<0.05. See also Supplemental data 2 and 3. In (c-d), the darker the color the higher the number of DEGs in a pathway, and the bigger the letters, the higher percentage of the pathway that is differential.

### Cytokinin affects the fungal cell cycle

Since we observed that CK inhibits sporulation, spore germination, and hyphal growth, and the cell cycle and DNA replication were down regulated by CK treatment in the transcriptome analysis, we examined the morphology and relative DNA quantity of *Bc* cells grown with and without CK (**Fig. 6**). We observed a significant reduction of ∼75% in cell size upon CK treatment, with cells also appearing rounder (**Fig. 6a,b,d**).The distance between the last two septa from the hyphal tip were reduced to about 30% of normal length (**Fig. 6e**). 8 hours after germination, CK grown hyphae produced less than half the number of septa when compared with mock grown hyphae (**Fig. 6f**), further confirming that cell replication is inhibited. Concurrently, we quantified Hoechst staining in mock and CK grown mycelia. Hoechst binds to DNA and has been used before to estimate DNA content in live cell nuclei (Gomes *et al*., 2018). CK grown cells were stained by Hoechst less than 50% of the amount of stain observed in mock grown cells (**Fig. 6c,g**). Images were taken under identical conditions. Reduction in Hoechst staining coupled with lower rates of cell replication and the transcriptomic data indicating downregulation of cell cycle and meiosis pathways, indicates that CK may be inhibiting mitosis in *Bc*.

**Fig. 6.**
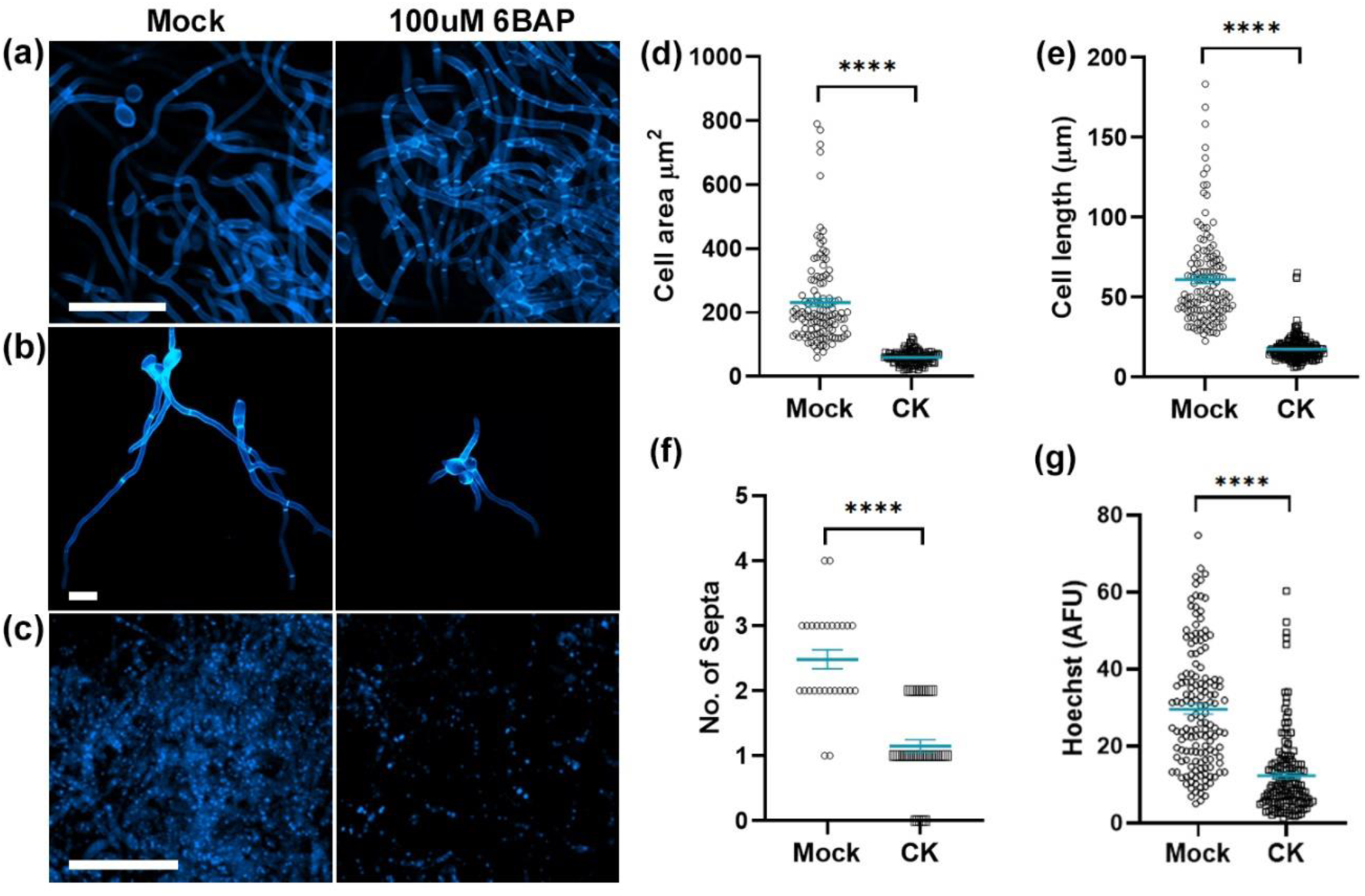
Cytokinin reduces *B. cinerea* cell elongation and DNA replication. *B. cinerea* was cultured in PDB liquid broth, with or without 100 µM CK (6-Benzylaminopurine). After 8 **(f)** or 16 hours **(a-e,g)**, growing hyphae were stained with calcofluor **(a,b,d,e,f)** or hoechst **(c,g)**, and imaged on an Olympus IX 81 confocal laser scanning microscope using a 405nm diode laser (1% power), under identical imaging conditions. **a-c:** bar=50 µM; **b:** bar=20 µM. Cell area **(d)**, distance between septa **(e)**, No. of septa in individual germinated hyphae **(f)** and DNA staining **(g)** were measured using Fiji-ImageJ. Graphs represent 3-6 biological repeats, N>120 (d,e), N>30 (f), N=170 (g). Statistically significant differences between Mock and CK samples were assessed using a two-tailed t-test, ****p<0.0001. Individual values graphed, blue bar represents mean ±SE.

### Cytokinin inhibits the fungal cytoskeleton

Our results demonstrate that CK inhibits fungal growth and development (**Figs. 1,2,5**). We hypothesized that CK affects a fundamental cellular process relevant to most fungi; a process that is crucial to execute the fast growth occurring in hyphal tips (Bartnicki-Garcia, 2002), growth that requires membrane remodeling (Riquelme *et al*., 2018). Based on the NGS results, we hypothesized that these affected processes, in addition to the cell cycle, are likely to be cytoskeletal integrity and/or cellular trafficking.

To examine cytoskeleton integrity, we first validated the expression levels of cytoskeletal genes shown to be differential in the transcriptomic data. These genes are listed in **Fig. 7a** (saturated blue color indicates downregulation), with the full expression data provided in **Supplementary Data 4**. We independently confirmed relative expression of 5 genes from the data set by qRT-PCR, selecting both down regulated and up-regulated genes from the transcriptome (**Fig.7b**). As such, we used the geometric mean of 3 housekeeping genes that are unrelated to the cytoskeleton for gene expression normalization. We transformed *B. cinerea* with the filamentous actin marker lifeactin-GFP (Schumacher, 2012), and proceeded to treat the transformed fungal cells with CK. We observed mislocalization of life-actin, which is normally localized to growing hyphal tips (Walker & Garrill; Berepiki *et al*., 2011), upon CK treatment. CK caused life-actin to be distributed more uniformly throughout the cells, and to lose most of its tip-specific localization (**Fig. 7c-d**). Analysis of corrected total fluorescence in mock and CK treated cells demonstrated that the ratio between life-actin in the tip of the cell, and the total cell, decreased greatly in the presence of CK (**Fig. 7d**). Note that the transformed fungus showed the characteristic “split tips” phenotype of lifeactin overexpression (Schumacher, 2012) in both mock and CK treated samples.

**Fig. 7.**
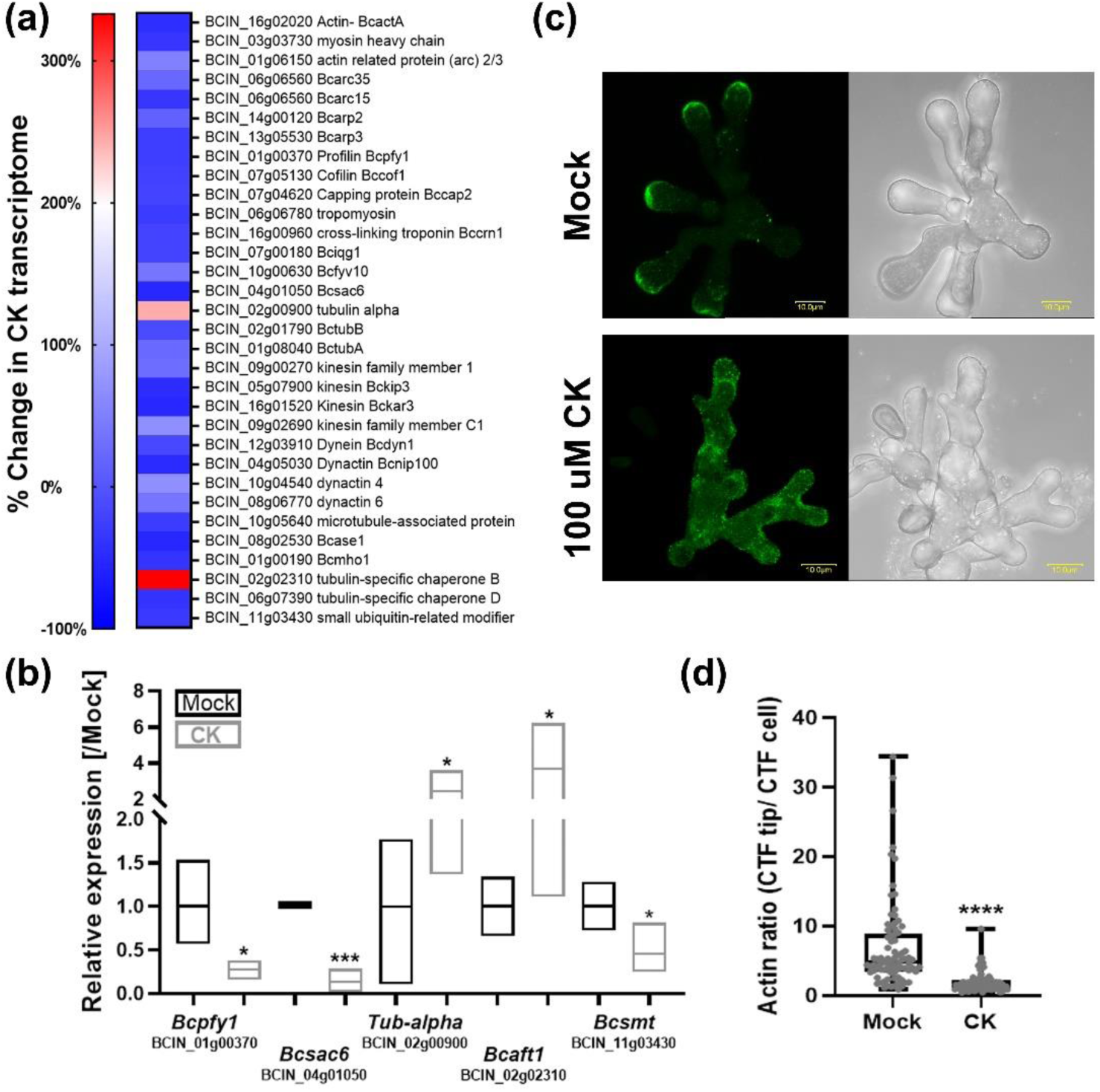
Cytokinin inhibits the fungal cytoskeleton. **(a)** Cytoskeleton related *B. cinerea* genes found to be differentially regulated in the RNAseq. See Supplemental data 4 for full list. **(b)** qRT-PCR validation of expression levels of 5 cytoskeleton related genes upon CK treatment. *B. cinerea* was grown in PDB with the addition of 100 µM CK (6-BAP) or without (Mock). Mock was set to 1. Gene expression values were normalized to a geometric mean of the expression of 3 housekeeping genes: Ubiquitin-conjugating enzyme E2, Iron-transport multicopper oxidase, and Adenosine deaminase. Floating bars represent minimum to maximum values of 3 biological repeats, line represents mean. Asterisks indicate significance in a two-tailed t-test with Welch’s correction, *p<0.05; ***p<0.001. **(c-d)** *B. cinerea* was transformed with lifeactin-GFP. Germinated spores were treated with Mock or CK and grown for 6 h prior to confocal visualization. **(c)** Representative images, bar=10 uM. **(d)** Analysis of corrected total fluorescence (CTF) of the ratio between life-actin at the tip of the cell and the total cell in Mock and CK treated cells. Three independent experiments were conducted with a total of 30 images analyzed, N>80 growing hyphae tips. Asterisks indicate significance in a two-tailed t-test with Welch’s correction, ****p<0.0001.

### Cytokinin inhibits fungal endocytosis

We and others (Marhavý *et al*., 2011; Gupta *et al*., 2020b) have previously shown that CK can influence cellular trafficking in plants. Further to our results demonstrating that CK inhibits the cytoskeleton in *B*c, and since the endocytic pathway was also found to be significantly downregulated by CK (**Fig. 5**), we examined the effect of CK on endocytosis in *Bc*. **Fig. 8** shows that 6-BAP inhibits endocytosis of the endocytic tracer FM-4-64, which is routinely used in fungi (Fischer-Parton *et al*., 2000), reducing the amount of endocytic vesicles by more than 50% (**Fig. 8a,b**). 6-BAP also caused a significant decrease in the size of the vesicles containing endocytic tracer (**Fig. 8c**), in both the 100 nM and 100 uM concentrations, similar to its effect on sporulation and spore germination (**Fig. 2**). This suggests that in parallel with the effect on the cytoskeleton, CK has a possible impact on membrane function and/ or fission of vesicles.

**Fig. 8.**
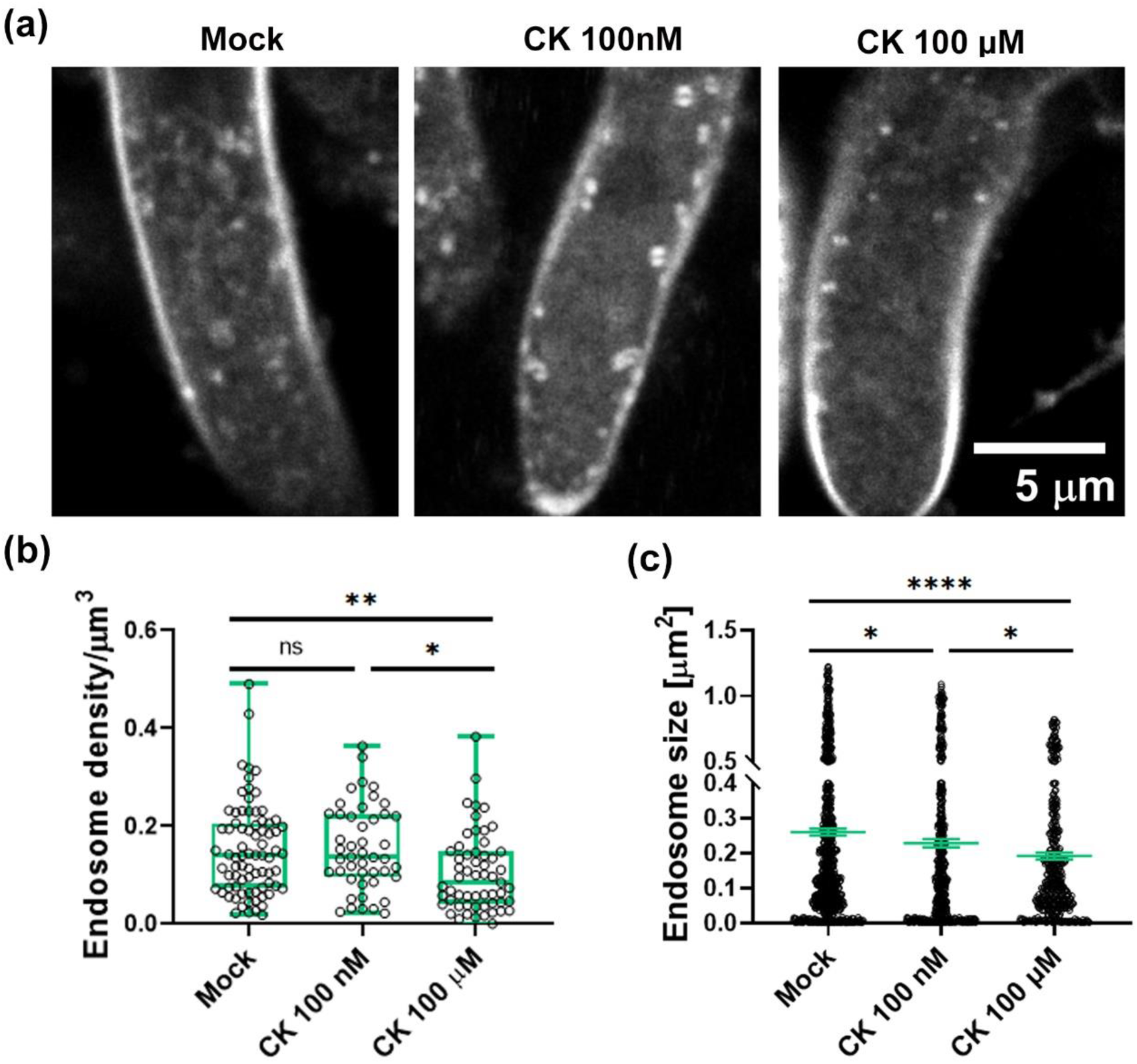
Cytokinin inhibits FM-4-64 endocytosis in growing *B. cinerea* hyphae. *B. cinerea* (*Bc*) was cultured in PDB liquid broth in the presence of 100 nM or 100 µM CK (6-Benzylaminiopurine) for 16 hours. **(a)** FM-4-64 endocytic vesicles in *Bc* hyphae. **(b)** quantification of the amount of endocytic vesicles; **(c)** quantification of the average size of vesicles. Measurements were done using the counting tool of Fiji. Quantification of results from 7 biological repeats. **b:** N>45, box-plot with all values displayed, box indicates inner-quartile ranges with line indicating median, whiskers indicate outer-quartile ranges. **c:** N>450, all values displayed, line indicates median ±SE. Asterisks indicate significance in one-way ANOVA with a Tukey post hoc test, *p<0.05; **p<0.01; ****p<0.0001; ns-non-significant.

#### Inhibition of endocytosis abolishes *B. cinerea* CK sensitivity

To further examine the effect of CK on endocytosis inhibition in *Bc*, we examined combined effects of CK and endocytosis inhibition using butanol treatments. 1-butanol inhibits phospolipases (Motes *et al*., 2005; Jia & Li, 2018), which have been linked to both growth and virulence in *Bc* (Schumacher *et al*., 2008) (see also **Fig. 3**), and was used as an endocytosis inhibitor in previous works in fungi (Boucrot *et al*., 2006; Sharfman *et al*., 2011; Li *et al*., 2012). We examined CK-mediated growth inhibition of both mycelia area and dry weight, in the presence or absence of the endocytosis inhibitor 1-butanol, as well as its structural analog 2-butanol, which does not inhibit endocytosis (Sharfman *et al*., 2011). **Fig. S4** demonstrates that upon endocytosis inhibition with 1-butanol, CK no longer affects fungal growth, while CK-mediated growth inhibition is retained in the mock and 2-butanol treatments. (**Fig. S4 a,b**). At 0.4%, 2-butanol did not inhibit fungal growth. At 0.8%, fungal growth inhibition was observed with 2-butanol, though to a significantly lesser degree than that observed with 1-butanol (**Fig. S4a**).

#### Inhibition of the cytoskeleton abolishes *B. cinerea* CK sensitivity

We further examined combined effects of CK and cytoskeleton disruption, using Benomyl (Ben) and Latrucnculin B (LatB). Benomyl depolymerizes microtubules, and has been previously used as a fungicide, and in studies of fungal cell cycle and cytoskeleton (Momany & Hamer, 1997; Peterson & Mitchison, 2002; Dub *et al*., 2013). Latrunculin B depolymerizes actin filaments, and has also previously been used for cytoskeletal studies in fungi (Ketelaar *et al*., 2012). We assayed the combined effect of CK and Ben or LatB on endocytosis, assaying endosome size and density. We observed that CK and Ben (**Fig. S5a,c**) or LatB (**Fig. S5b,d**) affect endocytic compartments in a similar manner, finding no enhancement of endocytosis inhibition when CK was combined with either drug (**Fig. S5**), suggesting that CK may inhibit endocytosis in part through its effect on the cytoskeleton, though down regulation of endocytic genes is also present (**Fig. 5**).

We also examined the combined effect of CK and Ben or LatB on growth inhibition of fungal mycelia. **Fig. S6** demonstrates that Ben and LatB mediated growth inhibition is not further enhanced by the addition of CK. Pursuant to the results presented in Figure 5, the size of hyphal cells treated with CK, Ben (**Fig. S6a**), LatB (**Fig. S6b**), or combinations of Ben/LatB and CK, was similarly reduced when compared with mock treated cells, demonstrating that CK may have a similar effect as Ben or LatB on the progression of the cell cycle. Additionally, colony mycelial area, which was reduced by Ben and LatB in a concentration dependent manner, remained mostly unaffected by the addition of CK in the background of these drugs (**Fig. S6c,d**).

### Cytokinin reduces yeast growth and endocytosis

Given our observations that CK can affect fundamental cellular processes in fungi, we examined inhibitory roles for CK in the growth of *Saccharomyces cerevisiae*, a budding yeast, and *Schizosaccharomyces pombe*, a fission yeast, the growth of which more closely resembles that of fungal hyphae. Growth curves were generated by measuring OD_600_ over time, as previously described (Laor *et al*., 2019). We observed that CK inhibits the growth of *S. cerevisiae* (**Fig. 9a,b**), and *S. pombe* (**Fig. S7a,b**) in a dose-dependent manner. We found that *S. pombe* was more strongly inhibited. Interestingly, trans zeatin was previously reported not to affect *S. pombe* cell division (Suzuki *et al*., 2001).

**Fig. 9.**
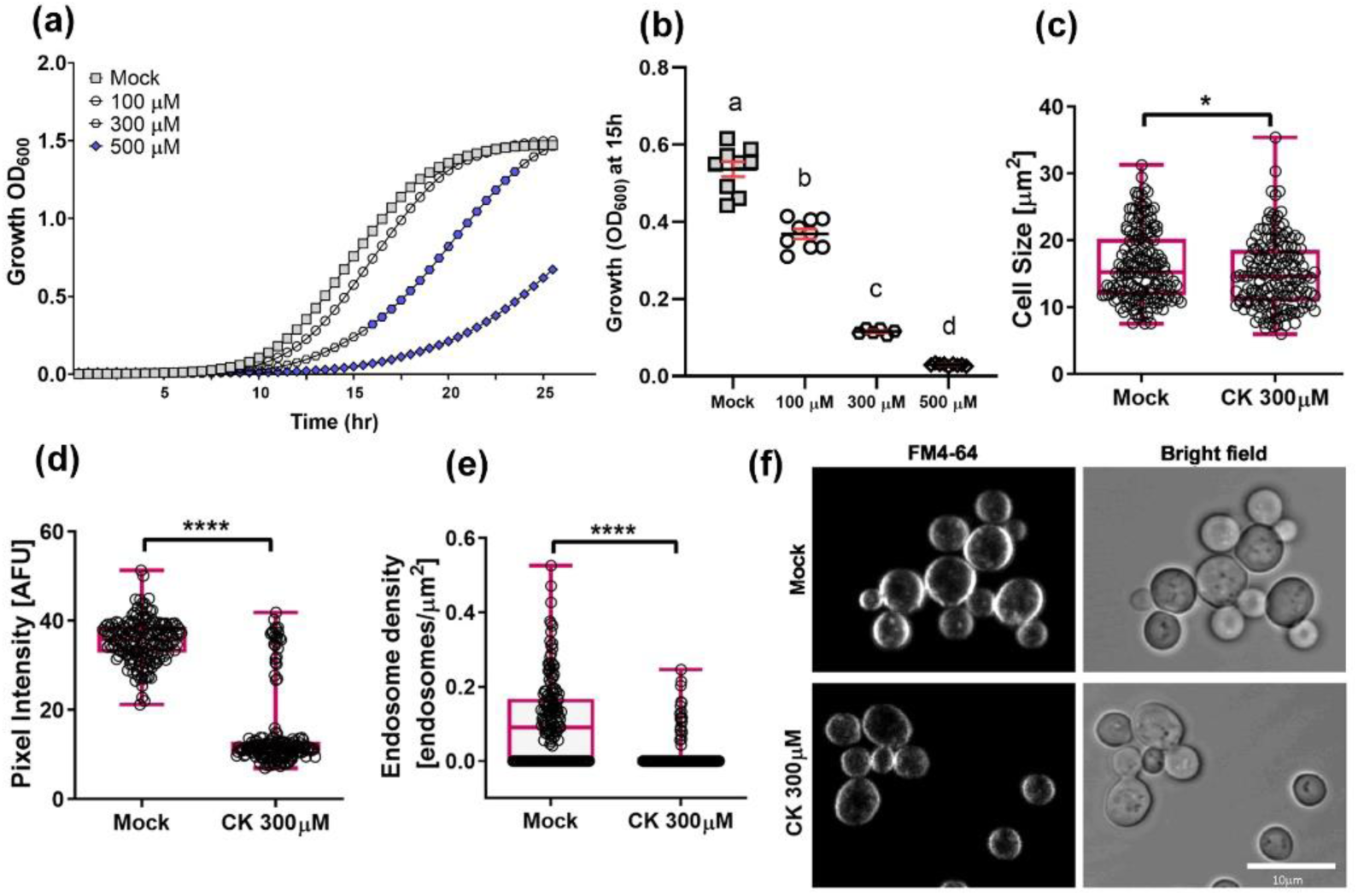
Cytokinin inhibits growth and endocytosis in budding yeast. **(a)** Wild-type *Saccharomyces cerevisiae* were grown over night at 30°C in minimal synthetic defined medium, treated with either 10uM NaOH (Mock) or with the addition of indicated concentrations of CK (6-Benzylaminopurine). Cells were incubated at 30°C for 25h, with continuous shaking. Average growth per time point for 3 experiments is presented, N=9. Blue color represents statistical significance from Mock treatment in a two-tailed t-test with Holm-Sidak correction, p<0.05. **(b)** Average growth (OD) at mid log phase (15h) in three independent experiments. Letters indicate significance in a one-way ANOVA with a post hoc Tukey test; p<0.0001. All points displayed; red lines indicate mean ±SE. **(c-f)** *S.cerevisiae* yeast cells were grown overnight at 30°C in YPD medium, diluted (OD_600_ = 0.2) and incubated for 6 hours in YPD media (Mock) or media supplemented with 300 µM CK (6-Benzylaminopurine). Cells were incubated with 24 μM FM4-64 (Invitrogen) at 4°C for 30 min. Subsequently, the FM4-64 containing medium was replaced with fresh medium and cultures was incubated at 28°C for 15 minutes. Confocal microscopy images were acquired using a Zeiss LSM780 confocal microscope. **(c)** cell size; **(d)** total internalized FM4-64 per cell represented by pixel intensity; **(e)** endosome density. **(f)** Representative images, Bar, 10 µm. Box-plots with all values displayed; line indicates median. **c-e:** N>160. Image analysis was performed using Fiji-ImageJ with raw images collected from 3 independent biological experiments. Endosome count measurements were done with the 3D Object counter tool and pixel intensity was measured using the measurement analysis tool. Asterisks represent statistical significance in a two-tailed t-test with a Mann-Whitney post hoc test, *p<0.05, ****p<0.0001.

To examine whether growth inhibition is mediated by endocytosis in yeast, as we found for *Bc*, we conducted endocytic assays in *S. cerevisiae* (**Fig. 9**) and *S. pombe* (**Fig. S8**). Yeast cultures were grown overnight, diluted to an OD_600_ of 0.2, and grown for a further six hours with or without CK. Cultures were then stained with FM4-64 (Vásquez-Soto *et al*., 2015). CK reduces cell size, internalization of FM4-64, endosome size, and endosome density in both *S. cerevisiae* (**Fig. 9c-f**) and *S. pombe* (**Fig. S8a-e**

In parallel, the effect of CK on the growth of *S. cerevisiae* endocytic mutants was examined. Genes known to be involved in endocytosis, the absence of which is known not to be lethal, were selected. *S. cerevisiae* knockout strains were generated as described in the methods section. Mutations in *YPT31*, a Rab/GTPase known to be involved in vesicular trafficking (Jedd *et al*., 1997), *SSA1*, known to be involved in clathrin vesicle uncoating (Krantz *et al*., 2013), *VPS1*, a dynamin-like GTPase known to be required for vacuolar sorting, cytoskeleton organization and endocytosis (Vater *et al*., 1992; Ekena *et al*., 1993) and *SPO14*, a phospholipase D protein required for sporulation (Rudge *et al*., 2002), were generated and examined. Three out of the four generated mutants, *ypt31Δ*, *ssa1Δ*, and *vps1Δ*, exhibited a partial rescue in CK mediated inhibition (**Fig. 10a-c,e,f**), growing significantly better in the presence of 300 and 500 µM 6-BAP than the wild type (WT) strain. The *spo14Δ* mutation did not rescue CK-mediated growth inhibition (**Fig. 10d-f).**

**Fig. 10.**
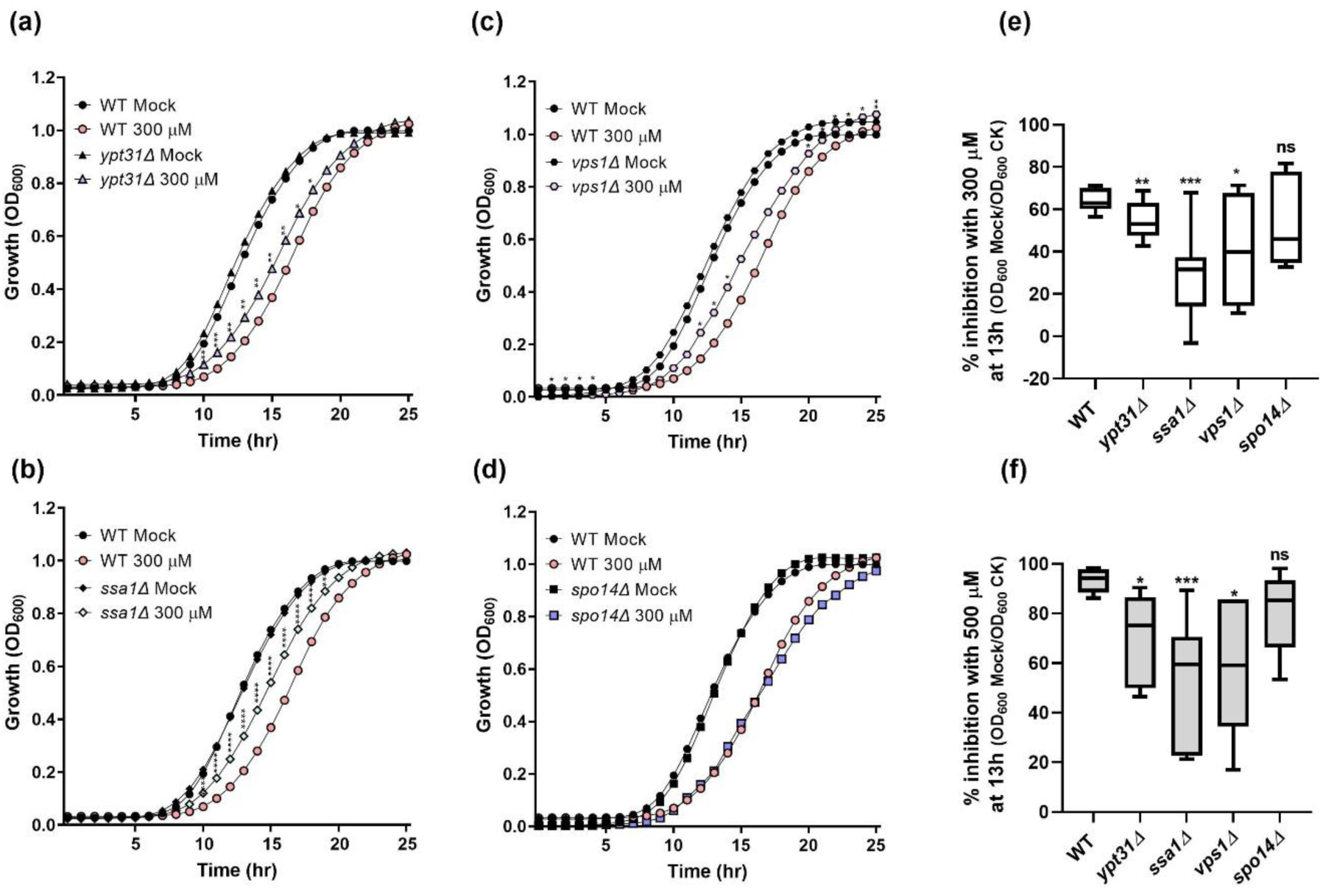
Cytokinin mediated growth inhibition is partially rescued in budding yeast endocytic mutants. *S. cerevisiae* Wild-type (WT, a-f), *ypt31Δ* (**a,e,f**)*, ssa1Δ* (**b,e,f**)*, vps1Δ* (**c,e,f**) and *spo14Δ* (**d,e,f**) were grown over night at 30°C for 25 h, in minimal synthetic defined medium, treated with either 10 µM NaOH (Mock) or with the addition of 300 µM (**a-e**) or 500 µM (**f**) CK (6-Benzylaminopurine). **(a-d)** Average growth per time point for three experiments is presented, N=9. Asterisks indicate statistical significance of each mutant with CK compared to WT with CK, in a two-tailed t-test with Holm-Sidak correction, *p<0.05, **p<0.01, ***p<0.001, ****p<0.0001. **(e-f)** Percentage of growth inhibition of each strain with 300µM CK (e) and 500 µM CK (f) as compared to mock treatment, at 13 h, in three independent experiments, N=9. Boxplots are shown with inter-quartile-ranges (box), medians (black line in box), and outer quartile whiskers, minimum to maximum values. Asterisks indicate significance in a one-way ANOVA with a post hoc Tukey test; *p<0.05; **p<0.01; ***p<0.001; ns (non-significant).

## Discussion

As reported previously by us and others, and as shown here (**Fig. 1**), CK promotes fungal disease resistance in plants (Choi *et al*., 2010; Gupta *et al*., 2020b). Direct effects of CK on fungal growth and development have not been investigated in depth in plant-free systems, and although a few studies have discussed some effects of CK on fungal pathogen growth (Babosha, 2009; Sharma *et al*., 2010), no mechanisms for CK antifungal activity have yet been reported. The present study was performed to examine the direct inhibitory effect of CK on different classes of fungal phytopathogens. We focused our efforts on three pathogens, *Bc, Sr* and *Fol*, with varied lifestyles and infection modes.

We found that CK treatment strongly inhibits *Bc* and *Sr*, and inhibits *Fol* more weakly (**Fig. 1d-j**). It has been demonstrated that pathogenic fungi of the *Fusarium* species complex are able to produce CKs endogenously, which might be the reason for its tolerance towards exogenous CK (Vrabka *et al*., 2018; Sørensen *et al*., 2018). Interestingly, varying levels of inhibitory activity were observed for CK in many phytopathogenic fungi (**Fig. S2**).

Examination of possible effects of CK on *Bc* development revealed that CK attenuates *Botrytis* sporulation, spore germination and germ tube elongation (**Fig. 2**), with differential effects of the CK concentration on different processes. Interestingly, genes involved in the regulation of growth, conidiation, germination, virulence, and pathogenicity in *Bc*, were strongly inhibited by CK treatment (**Fig. 3**). Several genes previously found to be involved in plant infection processes and defined as essential determinants for *Bc* pathogenicity (Zheng *et al*., 2000; Valette-Collet *et al*., 2003; Gourgues *et al*., 2004; Brito *et al*., 2006; Schumacher *et al*., 2008; Aguayo *et al*., 2011; Frías *et al*., 2019) were found to be significantly downregulated upon CK treatment (Figure 3c). This suggests that interference with the expression of virulence genes may also partly contribute to the inhibitory effects of CK against *Bc*. Inhibition of growth and spore germination together with downregulation of virulence genes in *Bc* could account for plant disease attenuation. This mechanism could complement the plant induced resistance mechanisms being activated by CK, which, as reported by us and others (Choi *et al*., 2010; Argueso *et al*., 2012; Gupta *et al*., 2020b), induces plant immunity even in the absence of a pathogen.

Damage in cell membrane integrity in fungi usually leads to the release of nucleic acids and proteins (Lewis & Papavizas, 1987; Ji *et al*., 2018; Wang *et al*., 2020). CK showed no effect on leakage of nucleic acids and proteins from *Bc* after treatment, and no change was observed in media conductivity of *Bc* after CK treatment, indicating that cell membrane permeability remained unchanged. Taken together, these results suggest that CK is not toxic to fungi. CK can thus be defined as possessing fungistatic, and not fungicidal, activity.

To understand the mode of action by which CK inhibited the growth of *Bc*, we conducted transcriptome profiling on *Bc* with and without CK. RNAseq data suggested that CK downregulates the cell cycle and cellular cytoskeleton and trafficking processes in *Bc*. Thus, we examined cell morphology and DNA replication after CK application. CK strongly reduced cell area, distance between septa, and likely, mitosis (**Fig. 6**). Cell division/septum formation is dependent on the signals generated during cell extension and growth, and nuclear division. Reduced cell growth and elongation effected by CK treatment correlates with the lesser number of septa in the treated cells. Hoechst staining revealed there is less DNA in CK treated cells, possibly due to inhibition of mitosis. Benomyl treatment, which is known to arrest the cell cycle, caused decreases in cell sizes that were not further augmented by CK (**Fig. S6**). Taken together, inhibition of cell division (septa formation) coupled with reduced DNA amounts and activity similar to benomyl, strongly support the notion that CK is inhibiting mitosis in fungal cells.

Also evident from the transcriptomic data was the effect of CK on the cytoskeleton and endocytic processes. Indeed, we found that CK caused mislocalization of actin at the growing tip of hyphae (**Fig. 7**), likely explaining, at least in part, the reduced hyphal growth observed (**Fig. 1**). CK inhibited the amount of endocytic vesicles in *Bc* (**Fig. 8**), *S. cerevisiae* (**Fig. 9**), and *S. pombe* (**Fig. S8**), indicating that it likely has an impact on membrane function and/ or fission of vesicles.

Cell elongation, in particular hyphal elongation, requires continuous addition of new plasma membrane, proteins, and cell wall material at the hyphal tip. Cellular trafficking and endo/exocytosis, which depen both on an intact cytoskeleton and on endocytic compartments, regulate the amount of membrane transferred towards this cellular growth, tightly controlling the amount of cellular material required for plasma membrane extension (Riquelme *et al*., 2018). Inhibition of the cytoskeleton and endocytosis by CK explains the reduced elongation and growth of *Bc* and yeast cells. Fission yeast may be more reliant on cellular trafficking for rapid cell elongation than budding yeast, and their growth is more similar to the growth of fungi than that of budding yeasts, possibly also explaining why fission yeast was more strongly inhibited by CK than budding yeast.

It has been previously suggested that certain classes of fungi possess CK receptors to be able to “sense plants”, a trait posited to have been required for land colonization by fungi (Hérivaux *et al*., 2017). We and others have previously demonstrated that plants activate CK signaling upon pathogen attack, and that CK serves as a cue to activate defense responses (Choi *et al*., 2010; Argueso *et al*., 2012; Gupta *et al*., 2020b). A possibility arising from our work is that when plants sense the presence of phytopathogenic fungi, an additional reason for the activation of CK pathways is to promote CK biosynthesis, thereby inhibiting the growth of the potential fungal attacker. We have shown here that CK inhibits growth in different types of fungi from different classes (basidiomycetes, ascomycetes) and lifestyles (soilborne, airborne, hemibiotroph, necrotroph), and yeasts (**Figs. 1, 2, 5, 8, S1, S2**), and cellular trafficking in *Bc*, *S. cerevisiae*, and *S. pombe* (**Figs. 6, 7, 9, S3**). Chemically or genetically attenuating cellular trafficking in *Bc* (**Fig. 7**), or *S. cerevisiae* (**Fig. 10**) respectively, resulted in loss of CK mediated growth inhibition. It would seem that, in employing CK as a fungal pathogen inhibitor, plants have succeeded in managing to target processes fundamental to growth, such that this inhibition is preserved all the way to budding yeast.

In the evolutionary war against plant pathogens, plants appear to have succeeded in creating a 1-2 mechanism for utilization of CK, previously believed to be only a “developmental” hormone. 1-The plant senses a pathogen and activates its CK response, leading to immunity signaling, which culminates in increased immunity and systemic pathogen resistance; and 2-the activation of CK response leads to the generation of increased CK levels, which inhibit the growth and development of fungal pathogens by targeting their cell cycle and trafficking machinery. Future work will validate the genetic targets of CK in fungi, and whether this fungistatic activity can be agriculturally adapted.

## Materials and Methods

### Pathogen growth conditions

*Botrytis cinerea* (strain Bcl16), *Sclerotium rolfsii* (Liarzi *et al*., 2020), *Fusarium oxysporum* f. sp. *lycopersici* (strain 4287) were cultivated on potato dextrose agar (PDA) at 22 ± 2 °C for *B. cinerea*, 26 ± 2 °C for *S. rolfsii* and 28 ± 2 °C for *F. oxysporum* for 5-7 days. Pathogen isolates were kindly gifted by Yigal Elad, David Ezra and Shay Covo.

### Chemical treatments

6-BAP (6-Benzylaminopurine), zeatin, kinetin, TDZ, (all in 10mM NaOH), adenine (1M HCL), Benomyl and latrunculinB (DMSO), 1-butanol and 2-butanol, were all obtained from Sigma-Aldrich, and added to media at indicated concentrations.

### Cytokinin level measurement

Cytokinin extraction was performed according to (Gupta *et al*., 2020b). Solvent gradients and MS-MS parameters are detailed in supplemental Table 1.

### Plant pathogenesis assays

*Bc* pathogenesis was performed as previously described (Gupta *et al*., 2020b).

*Fol* culture was grown in KNO_3_ media (yeast nitrogen base, sucrose, KNO_3_ and distilled water) at 28 °C for 5 days. Two week old tomato plants were treated with 100 µM 6-BAP (foliar spray) and inoculated with a spore suspension (10^6^ spores ml^-1^) (De Cal *et al*., 2000) using the root dip method (Wellman, 1939; Mes *et al*., 1999), 24 h later. The disease index (DI) was calculated after three weeks of using a 0–5 scale: 0, no symptoms; 1, ≤ 2% (healthy plant); 2, 3–30% (slight disease); 3, 31–60% (moderate disease); 4, 61–90% (severe disease), and 5, ≥ 91% (dead plant). For *Sr*, tomato plants were soil drenched with 100 µM of 6-BAP or mock (NaOH + Tween 20) one week prior to infection. After one week, 3-4 sclerotia of *Sr* were placed on the soil, ∼2cm from the plant stem. DI was assessed after 2 weeks.

### *B. cinerea* spore germination measurement

*B. cinerea* spores (10^6^ spores ml^-1^) were incubated in potato dextrose broth (PDB) containing 0 and 100 µM 6-BAP at 22 ± 2 °C for 8 hours. Conidia were washed twice in sterile water (1 ml), centrifuged at 12,000 rpm for 5 min, and re-suspended in 100 μL of sterile water. A 10 μL sample was analyzed under the microscope. The percentage of sporulation, spore germination and the length of germ tubes were measured using ImageJ software.

### Measurement of cellular leakage and electrolytes

*Bc* were treated with 6-BAP (0 and 100 µM) in PDB and incubated on a rotary shaker at 180 rpm for 24h at 22 °C. The mycelia were subsequently centrifuged and the aqueous solutions were used for measurement of the leakage of nucleic acids and proteins. The absorbance of the supernatant was measured at 595 nm using UV/VIS spectrophotometer. Leakage of proteins was quantified according to Bradford (Bradford, 1976). The release of nucleic acids in various treatments was measured by detecting the optical density at 260 nm. Electrical conductivity was measured using a conductivity meter (EUTECH instrument con510) after 24 h.

### *B. cinerea* qRT-PCR virulence assessment

*Bc* spores were grown in PDB with 0 and 100 µM 6-BAP in a rotary shaker at 180 rpm and 22 ± 2 ℃ for 24 hours. RNA was isolated using Tri-reagent (Sigma-Aldrich) according to the manufacturer’s instructions. cDNA was prepared and qRT-PCR was performed as previously described (Gupta *et al*., 2020b). The primer sequences for each gene, and primer pair efficiencies, are detailed in Supplementary Table 2. A geometric mean of the expression values of the three housekeeping genes: ubiquitin-conjugating enzyme E2 (ubce) (Silva-Moreno *et al*., 2016), Iron-transport multicopper oxidase, and Adenosine deaminase (Llanos *et al*., 2015) was used for normalization of gene expression levels. Relative gene expression levels were calculated using the 2^-△△Ct^ method (Pfaffl, 2001).

### RNA extraction, quality control and RNA-sequencing

Total RNA was extracted from liquid *B. cinerea* cultures grown for two days in 1/2 PDB with the addition of tobacco seedlings 5 days post germination (150 seedlings/ 50 mL media), mock or supplemented with 25uM 6BAP, 50 mg fungal mass per sample, using the Norgen total RNA purification kit (Norgen Biotek corp.) according to the manufacturer’s instructions. RNA yield and purity was measured by Nanodrop (ND-1000 Spectrophotometer, Wilmington, USA), and validated by Bioanalyzer 2200 TapeStation (Agilent Technologies, California, USA). cDNA libraries with multiplexing barcodes were prepared using the TrueSeq RNA kit (Illumina, San Diego, CA, USA). Libraries were evaluated with Qbit and TapeStation (Agilent Technologies, California, USA). Pooled libraries of the 6 samples were sequenced on one lane of an Illumina Hiseq 2500 instrument using a 60-bp single-end RNA-Seq protocol, to obtain ∼20 million reads per sample. Sequencing was performed at the Weizmann Institute of Science, Israel.

### Transcriptome analysis

Raw-reads were subjected to a filtering and cleaning procedure. The Trimmomatic tool was used to filter out adapter sequences, remove low quality sequences by scanning a 4-base wide sliding window, cutting when the average quality per base drops below <15 and finally, removal of reads shorter than 36 bases (Bolger *et al*., 2014). Clean reads were mapped to the reference genomes of *B. cinerea* B05.10 (assembly accession GCF_000143535.2) (Staats & van Kan, 2012) using STAR software (Dobin *et al*., 2013). Gene abundance estimation was performed using Cufflinks version 2.2 (Trapnell *et al*., 2012) combined with gene annotations from genbank. Heatmap visualization was performed using R Bioconductor (Gentleman *et al*., 2004). Gene expression values were computed as FPKM. Differential expression analysis was completed using the DESeq2 R package (Love *et al*., 2014). Genes with an adjusted p-value of no more than 0.05 and log2FC greater than 1 or lesser than -1 were considered differentially expressed. PCA was calculated using the R function prcomp. We submitted the raw sequencing data generated in this study to NCBI under bioproject accession number PRJNA718329.

The gene sequences were used as a query term for a search of the NCBI non-redundant (nr) protein database that was carried out with the DIAMOND program (Buchfink *et al*., 2014). The search results were imported into Blast2GO version 4.0 (Conesa *et al*., 2005) for gene ontology (GO) assignments. Gene ontology enrichment analysis was carried out using Blast2GO program based on Fisher’s Exact Test with multiple testing correction of false discovery rate (FDR). KOBAS 3.0 tool (http://kobas.cbi.pku.edu.cn/kobas3/?t=1) (Xie *et al*., 2011) was used to detect the statistical enrichment of differential expression genes in KEGG pathway and Gene Ontology (GO).

### Cell elongation and DNA content

*Bc* spores were cultured in PDB with or without 6-BAP (100 µM) at 22 °C for 8 (septa counting in individual hyphae) or 16 h. The cells were then collected and stained with 1 g l^−1^ calcofluor white M2R at room temperature for 15 min, or 15 ug ml^−1^ Hoechst 33342 (Sigma-Aldrich) in the dark for 1 h in a humid chamber at room temperature (Dub *et al*., 2013).

Nuclei and septa were visualized with an Olympus 398 IX 81 confocal microscope (Fluoview 500) equipped with a 60× 1.0 NA PlanApo water immersion objective. Images were acquired using a 399 nm excitation laser (1% power), with emission collected in the range 385-420 nm. Images were analyzed using Fiji-ImageJ. Cell size and length (septal distance) were measured using the area measurement tool. Septa were counted using the counter tool. DNA staining was assessed using mean intensity measurement.

### *B. cinerea* transformation

We used the plasmid pNDH-OLGG (Schumacher, 2012) to generate *B. cinerea* expressing lifeact-GFP. The transformation cassette was amplified using primers GA 34F/34R (Supplementary Table 4). We transformed *B. cinerea* using PEG mediated transformation. 0.125% lysing enzyme from Trichoderma harzianum (Sigma–Aldrich, Germany) was used for protoplast generation. Following transformation, protoplasts were plated on SH medium (sucrose, Tris-Cl, (NH_4_)_2_HPO_4_ and 35µg/ ml hygromycin B). Colonies that grew after 2 days were transferred to PDA-hygromycin, and conidia were re-spread on selection plates to obtain a monoconidial culture. Transformants were visualized under a confocal microscope and screened with primers GA 44F/44R and GA 31F/31R (Supplementary Table 4). Confirmed transformants were stored at 80°C and used for further experiments. Confocal microscopy images were acquired using an Olympus 398 IX 81 confocal microscope (Fluoview 500) equipped with a 60×1.0 NA PlanApo water immersion objective. eGFP images were acquired using a 488 nm excitation laser (1% power), with emission collected in the range 488-509 nm. Corrected total fluorescence was quantified using ImageJ.

### *B. cinerea* endocytosis

*Bc* hyphae were grown in PDB with or without 6-BAP (100nM, 100 µM), Benomyl (sigma, cat no. 17804-35-2), or LatrunculinB (sigma, cat no. 76343-94-7) for 16 h at 22 °C, after which the cells were collected and stained with 5 µM FM4-64 on a glass coverslip. We acquired confocal microscopy images using a Zeiss LSM780 confocal microscope equipped with a 63×/1.15 Corr Objective. FM-4-64 images were acquired with a 514 nm excitation laser (4% power), with the emission collected in the range of 592–768 nm. Images of 8 bits and 1024 × 1024 pixels were acquired using a pixel dwell time of 1.27, pixel averaging of 4, and pinhole of 1 airy unit (1.3 µM). Image analysis (18-24 images per treatment collected in three independent experiments) was conducted with Fiji-ImageJ using the raw images and the 3D object counter tool and measurement analysis tool (Schindelin *et al*., 2012). Endosome density and size were calculated automatically by the software tool considering 1.3 µM depth, based on single optical sections. **Budding (*Saccharomyces cerevisiae*) and fission (*Schizosaccharomyces pombe*) yeast growth** Wild-type haploid yeast strains were grown over night at 30°C. *S. cerevisiae* cells were grown in synthetic defined (SD) medium and *S. pombe* cells were grown in Edinburgh minimal medium (EMM), without (mock) or with the addition of indicated concentrations of CK (6-BAP). *S. cerevisiae* and *S. pombe* were diluted to an OD_600_ of 0.01. 200 µL of cells were plated in 96 well plates and incubated at 30°C for 25h (*S. cerevisiae*) or 45h (*S. pombe*), with continuous shaking. OD_600_ was measured using a TecanTM SPARK 10M plate reader. Experiment was repeated three times with similar results. WT strains of *S. cerevisiae* and *S. pombe* were kind gifts from Martin Kupiec and Ronit Weisman.

### Budding (*S. cerevisiae*) and fission (*S. pombe*) yeast endocytosis

*Saccharomyces cerevisiae* was grown overnight in YPD media, then the culture was diluted (OD_600_ = 0.2) and incubated for 6 hours in YPD (mock) or YPD supplemented with 300 µM 6-BAP. *Saccharomyces pombe* was grown overnight in YE. The cultures were then diluted (OD_600_ = 0.2) and incubated for 6 hours in YE media (mock) or media supplemented with 100 µM 6-BAP. Cell cultures were collected by centrifugation at 5,000 rpm for 4 minutes, and re-suspended in fresh growth medium. FM4-64 staining was performed as described (Vásquez-Soto *et al*., 2015). Cells were incubated with 24 μM FM4-64 (Invitrogen) for 30 min at 4°C. Subsequently, the FM4-64 containing medium was replaced with fresh medium, and cultures was incubated for 15 minutes at 28°C. To observe FM4-64 distribution, 5 µl of the suspensions were placed on a slide and live confocal imaging was performed. Confocal microscopy images were acquired using a Zeiss LSM780 confocal microscope equipped with Objective LD C-Apochromat 63×/1.15 Corr. Acquisition settings were designed using an excitation laser wavelength of 514 nm (4% power). The emission was then collected in the range of 592–768 nm. Images of 8 bits and 1024 × 1024 were acquired using a pixel dwell time of 1.27, pixel averaging of 4 and pinhole of 1 airy unit. Bright field was acquired using the T-PMT (transmitted light detector). Image analysis was performed using Fiji-ImageJ with the raw images (Schindelin *et al*., 2012), endosome count and size measurements were performed with the 3D Object counter tool and pixel intensity was measured using the measurement analysis tool.

### Construction of budding yeast mutant strains

The *YPT31*, *SSA1, VPS1 and SPO14* genes were disrupted in wild-type yeast strain (BY4741) via homologous recombination using PCR fragments amplified from the plasmid pFA6a-KanMX6 as a template with suitable primers. Gene replacement was validated by PCR with suitable primers. Primer sequences are provided in Supplemental Table 3.

### Data analysis

All data is presented as the average ±SEM. We analyzed the statistical significance of differences between two groups using a two-tailed t-test, with additional post hoc correction where appropriate, such as FDR calculation with Holm-Sidak correction, and Welch’s correction for t-tests between samples with unequal variance. We analyzed the statistical significance of differences among three or more groups using analysis of variance (ANOVA). Regular ANOVA was used for groups with equal variances, and Welch’s ANOVA for groups with unequal variances. Significance in differences between the means of different samples in a group of 3 or more samples was assessed using a post-hoc test. The Tukey post-hoc test was used for samples with equal variances, when the mean of each sample was compared to the mean of every other sample. The Bonferroni post-hoc test was used for samples with equal variances, when the mean of each sample was compared to the mean of a control sample. The Dunnett post-hoc test was used for samples with unequal variances. Statistical analyses were conducted using Prism8^TM^.

## Acknowledgements

The authors would like to thank Yigal Elad for the *B. cinerea* strain (Bcl-16), Shay Covo for the *F. oxysporum* f. sp. *lycopersici* strain (4287), David Ezra for gifting various fungal pathogen isolates, and Martin Kupiec and Ronit Weisman for WT strains of *S. cerevisiae* and *S. pombe*. This work was supported by the Israel Science Foundation grant No. 1759/20 to MB. GA is supported by the Indo-China ARO Postdoctoral Fellowship Program. MB thanks members of the Bar group for continuous discussion and support.

## Author contributions

Conceptualization: MB, RG. Design: RG, LP, GA, DL, TY, MB. Methodology & experimentation: RG, LP, GA, DL, NK. Analysis: RG, LP, GA, DL, NS, UG, MB. Manuscript: RG, LP, GA, DL, UG, MB.

**The authors declare no competing interest.**

## Data availability statement

The authors declare that the data supporting the findings of this study are available within the paper and its supplementary information files. Raw data is available from the corresponding author upon reasonable request.

## Supplemental Information

**Fig. S1.**
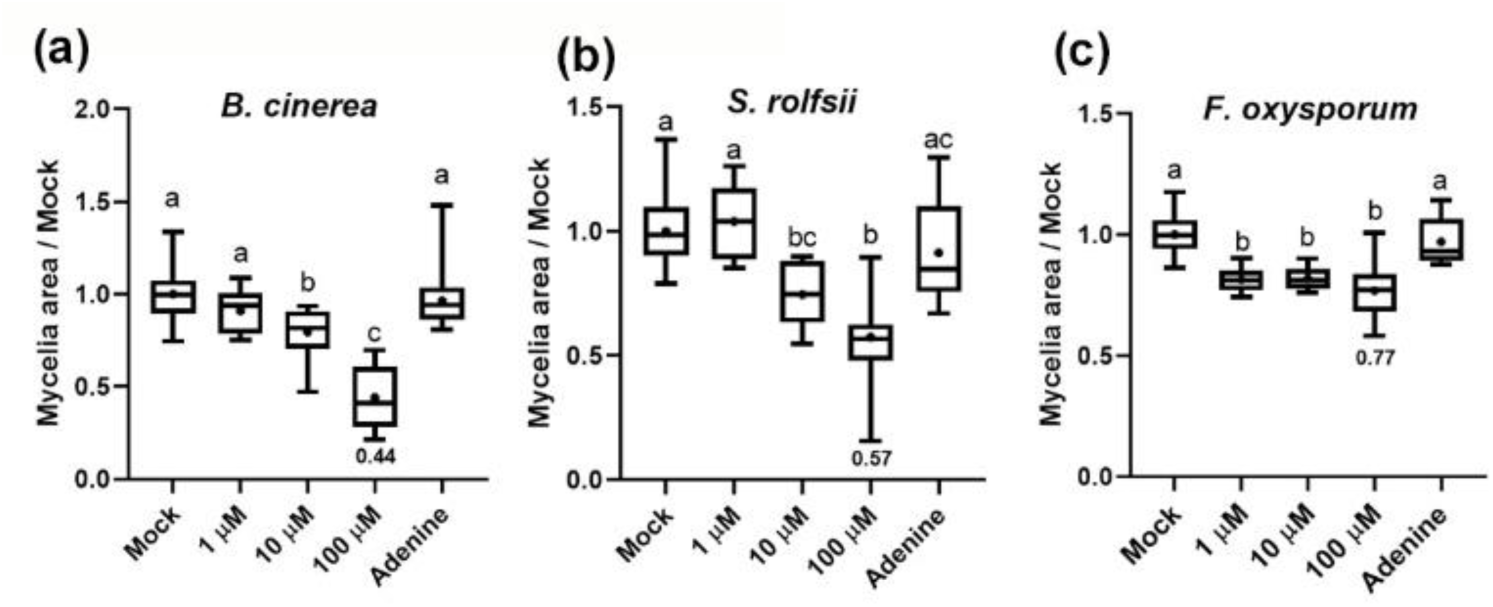
Direct effect of cytokinin on mycelial growth of phytopathogenic fungi: 6-BAP dose response. Dose response of *B. cinerea* **(a)**, *S. rolfsii* **(b)** and *F. oxysporum* **(c)** to CK-different concentrations of 6-Benzylaminopurine as indicated. Graphs represent 3 biological repeats ±SE, N>6. Letters indicate significance in a one-way ANOVA, ****p<0.0001 in all cases, with a Tukey post-hoc test. Box-plot displays minimum to maximum values, with inner quartile ranges indicated by box and outer-quartile ranges by whiskers. Line indicates median, dot indicates mean.

**Fig. S2.**
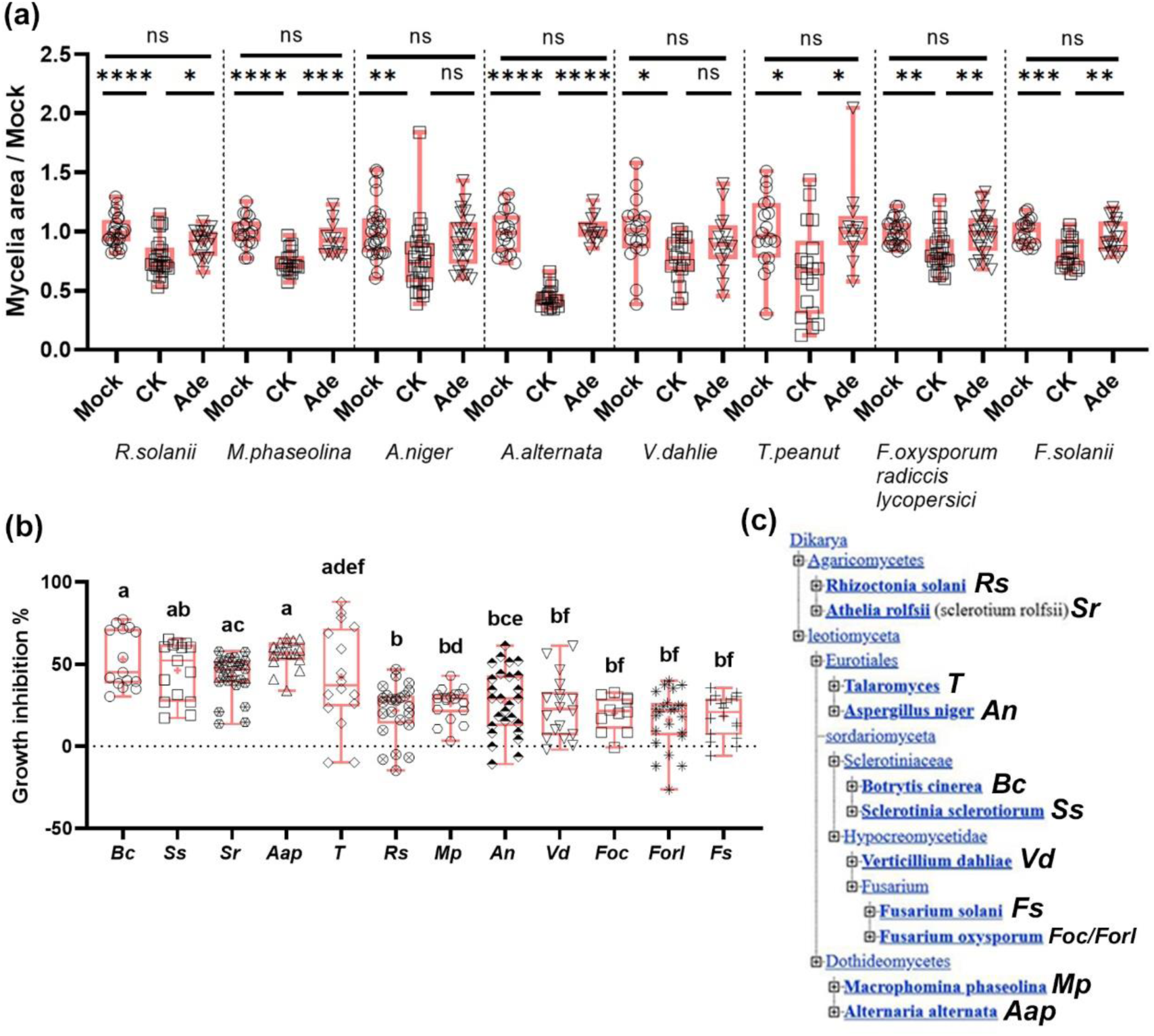
Cytokinin inhibits fungal growth. **(a)** Growth of different fungi cultured on potato dextrose agar (PDA) plates in the presence of 100 µM CK (6-Benzylaminopurine), or the control Adenine (Ade). Quantification of results from 4-6 biological repeats ±SE, N>12. Asterisks indicate significance in one-way ANOVA with a Tukey post hoc test, *p<0.05, **p<0.01, ***p<0.001; ****p<0.0001; ns (non-significant). **(b)** Comparison of the inhibition level of CK for different fungi. Letters indicate significance in a two-tailed t-test, p<0.004. The phylogeny is detailed in **(c). a-b:** Box plots with all individual values shown, line indicates median.

**Fig. S3.**
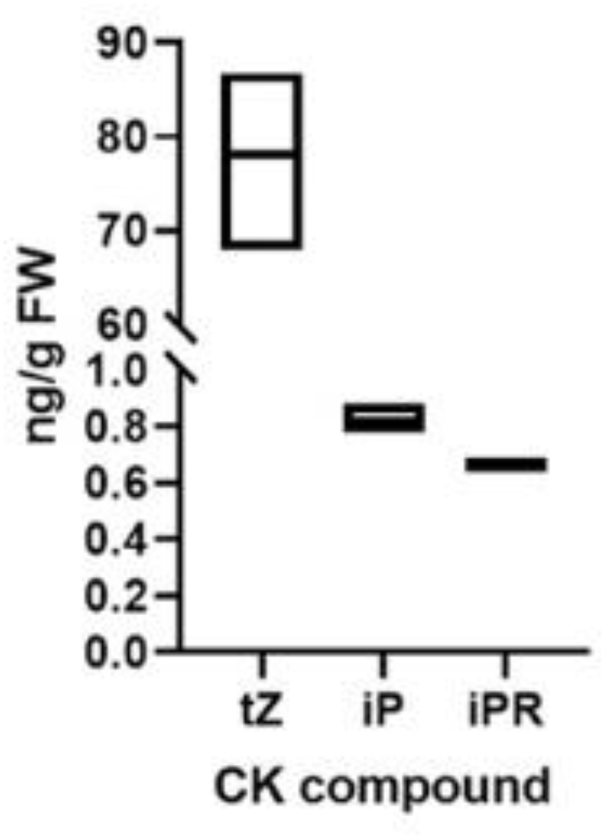
Quantification of active CKs in tomato leaves using LC–MS-MS. Fresh ground tissue powder was iso-propanol/methanol extracted in the presence of deuterium-labelled internal standards. LC–MS-MS analyses were conducted using a UPLC-Triple Quadrupole MS (WatersXevo TQMS). Acquisition of LC–MS data was performed using Mass Lynx V4.1 software (Waters). Quantification of CKs (tZ, trans zeatin, iP, isopentenyladenine and iPR, iP riboside) was done using isotope-labeled internal standards (IS), as described in the methods section. Quantification of results from 3 biological replicas ±SE. Bars represent minimum-maximum values range, with line indicating mean.

**Fig. S4.**
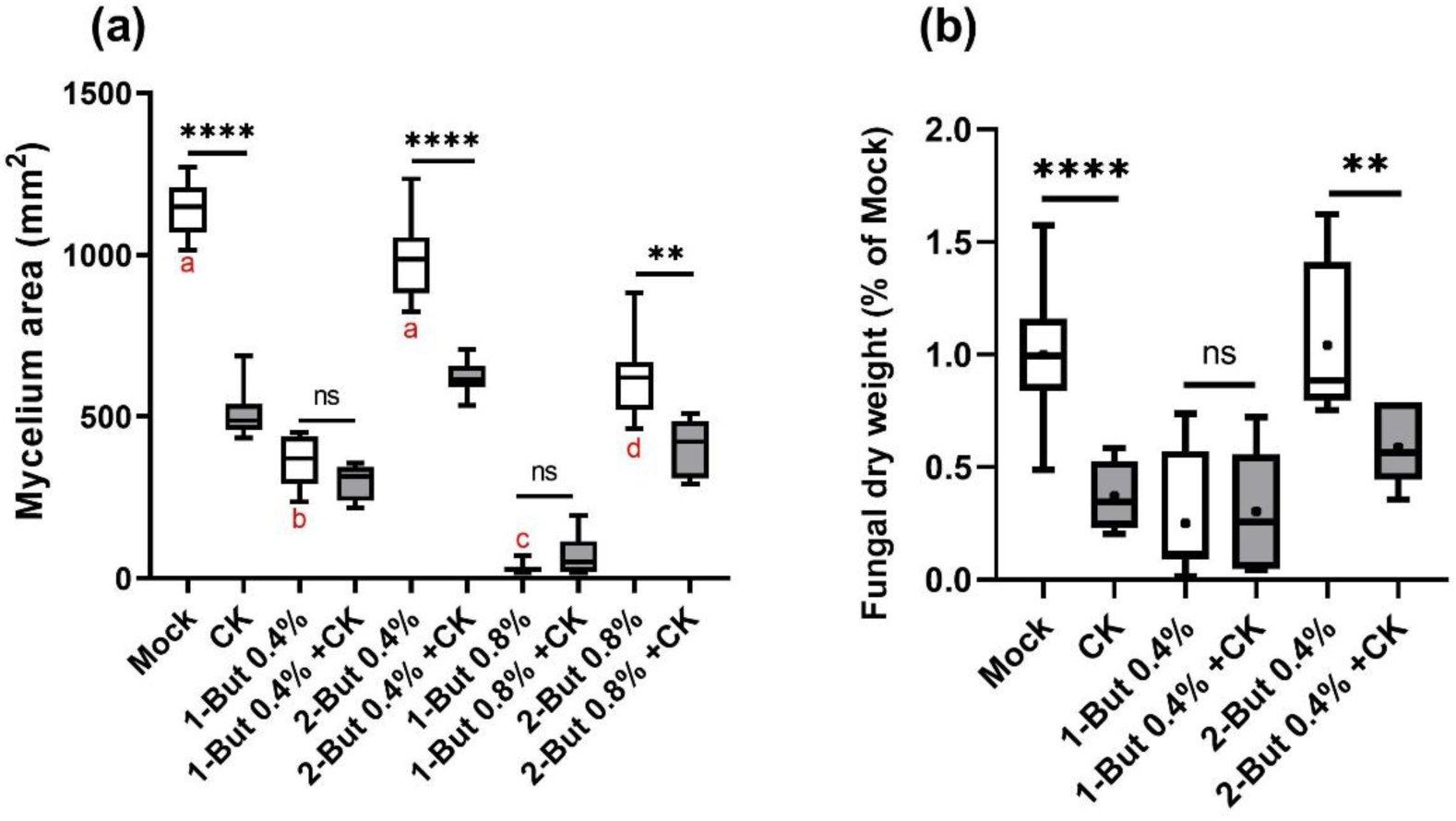
Inhibition of endocytosis abolishes *B. cinerea* cytokinin sensitivity. *B. cinerea* was cultured on 1/2 PDA plates (a) or in PDB liquid broth (b) in the presence or absence of 100 µM CK (6-Benzylaminopurine), the endocytosis inhibitor 1-butanol (v/v%), or control compound 2-butanol (v/v%) for 48 (a) or 96 (b) hours. **(a)** Mycelium area; measurements were done using Fiji. **(b)** Dry weight %. Quantification of results from 4 biological repeats ±SE, a: N>6, b: N>9. Boxplots are shown with inter-quartile-ranges (box), medians (black line in box), means (dot) and outer quartile whiskers, minimum to maximum values. Asterisks indicate significance in one-way ANOVA with a Dunnett post hoc test, **p<0.01; ****p<0.0001; ns (non-significant). Red letters in (a) indicate significance between the different treatments without CK in a one-way ANOVA with a Dunnett post hoc test, p<0.01.

**Fig. S5.**
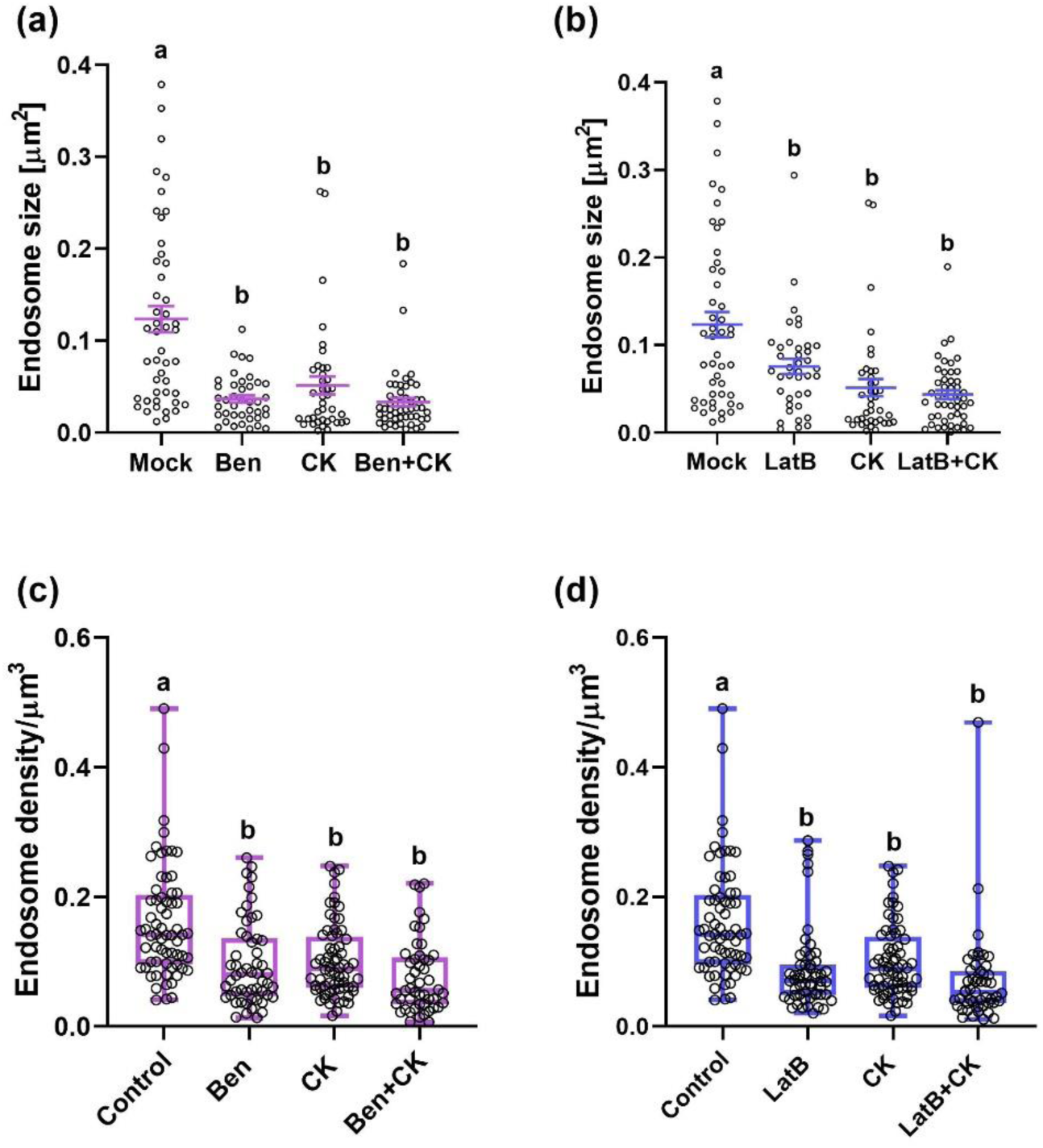
Inhibition of the cellular cytoskeleton abolishes *B. cinerea* cytokinin sensitivity-cellular trafficking. *B. cinerea* (*Bc*) was cultured in PDB liquid broth in the presence of 100 µM CK (6-Benzylaminiopurine) and/ or 1 uM Benomyl (Ben; **a,c**) or Latrunculin B (LatB; **b,d**) for 8 hours. FM-4-64 endocytic vesicles were analyzed in growing hyphae. Measurements were done using the counting tool of Fiji. **(a,b)** Quantification of the average size of vesicles from 3 biological repeats, N>40 images, the average endosome size per image was used for the analysis. All points displayed, mean ±SE is indicated. **(c,d)** Quantification of the amount of endocytic vesicles from 5 biological repeats, N>50 images. Box-plots with all values displayed, box indicates inner-quartile ranges with line indicating median, whiskers indicate outer-quartile ranges. Different letters indicate significance between samples in a one-way ANOVA with a Dunnett post hoc test, a,b: p<0.0059; c,d: p<0.0001.

**Fig. S6.**
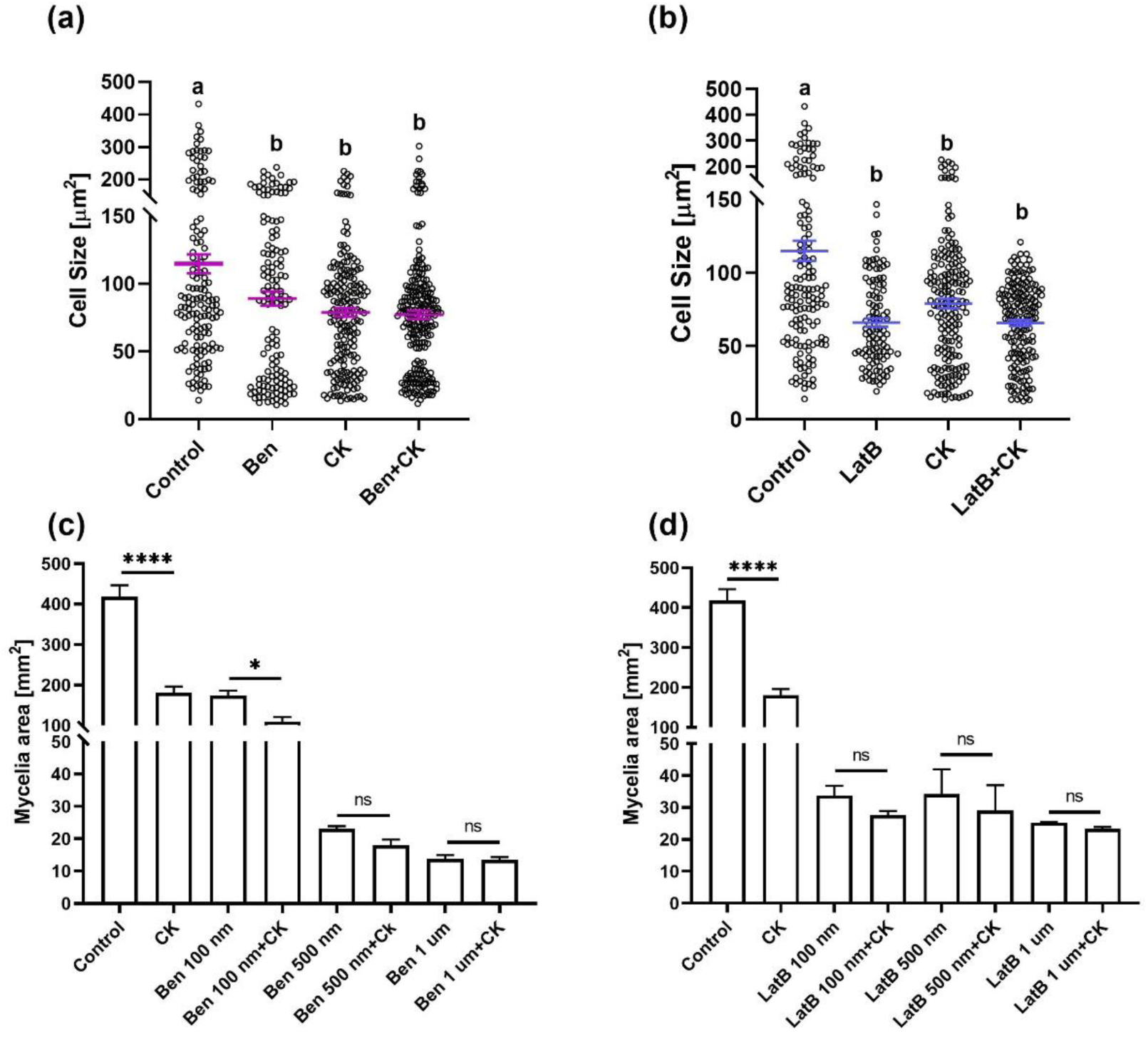
Inhibition of the cellular cytoskeleton abolishes *B. cinerea* cytokinin sensitivity-cell cycle and growth. **(a-b)** *B. cinerea* (*Bc*) was cultured in PDB liquid broth in the presence of 100 µM CK (6-Benzylaminiopurine) and/ or 1 uM Benomyl (Ben; **a**) or Latrunculin B (LatB; **b**) for 8 hours. Cell size was quantified in 3 experiments, N>100. All points displayed, mean ±SE is indicated. Different letters indicate significance between samples in a one-way ANOVA with a Dunnett post hoc test, p<0.021. **(c-d)** *B. cinerea* (*Bc*) was cultured on PDA plates in the presence of 100 µM CK (6-Benzylaminiopurine) and/ or indicated concentrations of Benomyl (Ben; **c**) or Latrunculin B (LatB; **d**) for 48 hours. Graph represents mean ±SE, N=3. Asterisks indicate significance between samples in a one-way ANOVA with a Bonferroni post hoc test, *p<0.05; ****p<0.0001; ns=not significant.

**Fig. S7.**
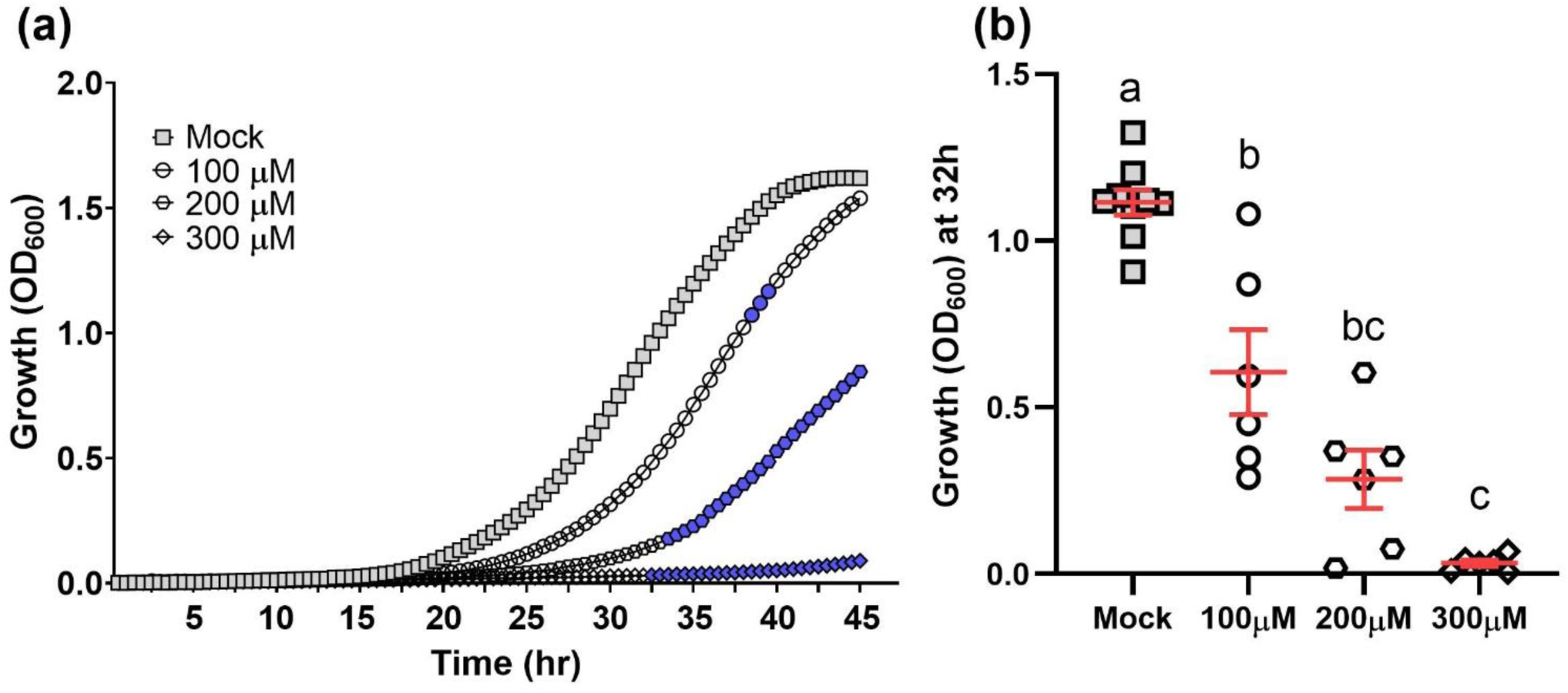
Cytokinin inhibits the growth of fission yeast. **(a)** Wild-type *Schizosaccharomyces pombe* cells were grown over night at 30°C in minimal EMM medium, treated with either 10uM NaOH (Mock) or with the addition of indicated concentrations of CK (6-Benzylaminopurine). Cells were incubated at 30°C for 45h, with continuous shaking. Average growth per time point for three independent experiments is presented, N=6. Blue color represents statistical significance in a two-tailed t-test with Holm-Sidak correction, p<0.05. **(b)** Average growth (OD) at mid log phase (32h) in three independent experiments. Letters indicate significance in a one-way ANOVA with a post hoc Tukey test; p<0.0001. All points displayed; red lines indicate mean ±SE.

**Fig. S8.**
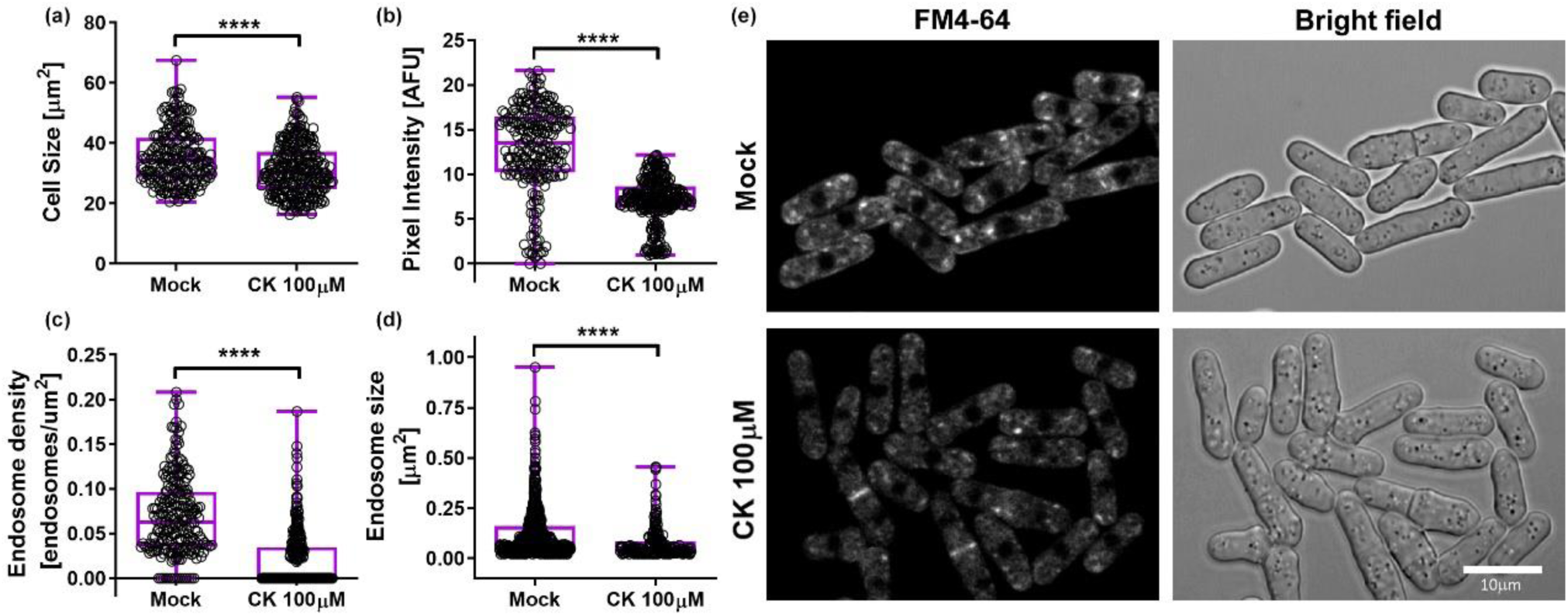
Cytokinin inhibits FM-4-64 endocytosis in fission yeast. *S. pombe* were grown overnight at 30°C in YE medium, diluted (OD_600_ = 0.2) and incubated for 6 hours in YE medium (Mock) or medium supplemented with 100 µM CK (6-Benzylaminopurine). Cells were incubated with 24 μM FM4-64 (Invitrogen) at 4°C for 30 min. Subsequently, the FM4-64 containing medium was replaced with fresh medium and cultures was incubated at 28°C for 15 minutes. Confocal microscopy images were acquired using a Zeiss LSM780 confocal microscope system with Objective LD C-Apochromat 63×/1.15 Corr. Acquisition settings were designed using an excitation laser wavelength of 514 nm (4% power). The emission was then collected in the range of 592–768 nm. Bright field was acquired using the T-PMT (transmitted light detector). **(a)** cell size; **(b)** total internalized FM4-64 per cell represented by pixel intensity; **(c)** endosome density; **(d)** endosome size. **(e)** Representative images, Bar, 10 µm. Box-plots with all values displayed; line indicates median. **a-c:** N>220; **d:** N>180. Image analysis was performed using Fiji-ImageJ with raw images collected from 3 independent biological experiments. Endosome count and size measurements were done with the 3D Object counter tool and pixel intensity was measured using the measurement analysis tool. Asterisks represent statistical significance in a two-tailed t-test with a Mann-Whitney post hoc test, ****p<0.0001.

**Supplementary Table 1.** Solvent gradients and MS-MS parameters for CK quantification.

Solvent gradient program for cytokinins:
| Time (min) | Phase A % | Phase B % |
| --- | --- | --- |
| Initial | 95 | 5 |
| 0.5 | 95 | 5 |
| 14 | 50 | 50 |
| 15 | 5 | 95 |
| 18 | 5 | 95 |
| 19 | 95 | 5 |
| 22 | 95 | 5 |

LC-MS-MS parameters for quantifications:
| Analite<br>and IS | Retention time<br>(min) | Ionization<br>Mode | MRM<br>transition<br>(m/z) | Dwell<br>time<br>(msec) | Cone<br>(V) | Collision<br>(V) |
| --- | --- | --- | --- | --- | --- | --- |
| t-Z | 2.34 | positive | 220>202<br>220>136 | 78 | 26 | 14<br>18 |
| <sup>2</sup>H5-t-Z | 2.35 | positive | 225>137<br>225>207 | 78 | 22 | 18<br>12 |
| iP | 4.86 | positive | 204>136<br>204>69 | 30 | 20 | 14<br>16 |
| <sup>2</sup>H6-iP | 4.82 | positive | 210>137<br>210>75 | 30 | 24 | 16<br>18 |

**Supplementary Table 2.**
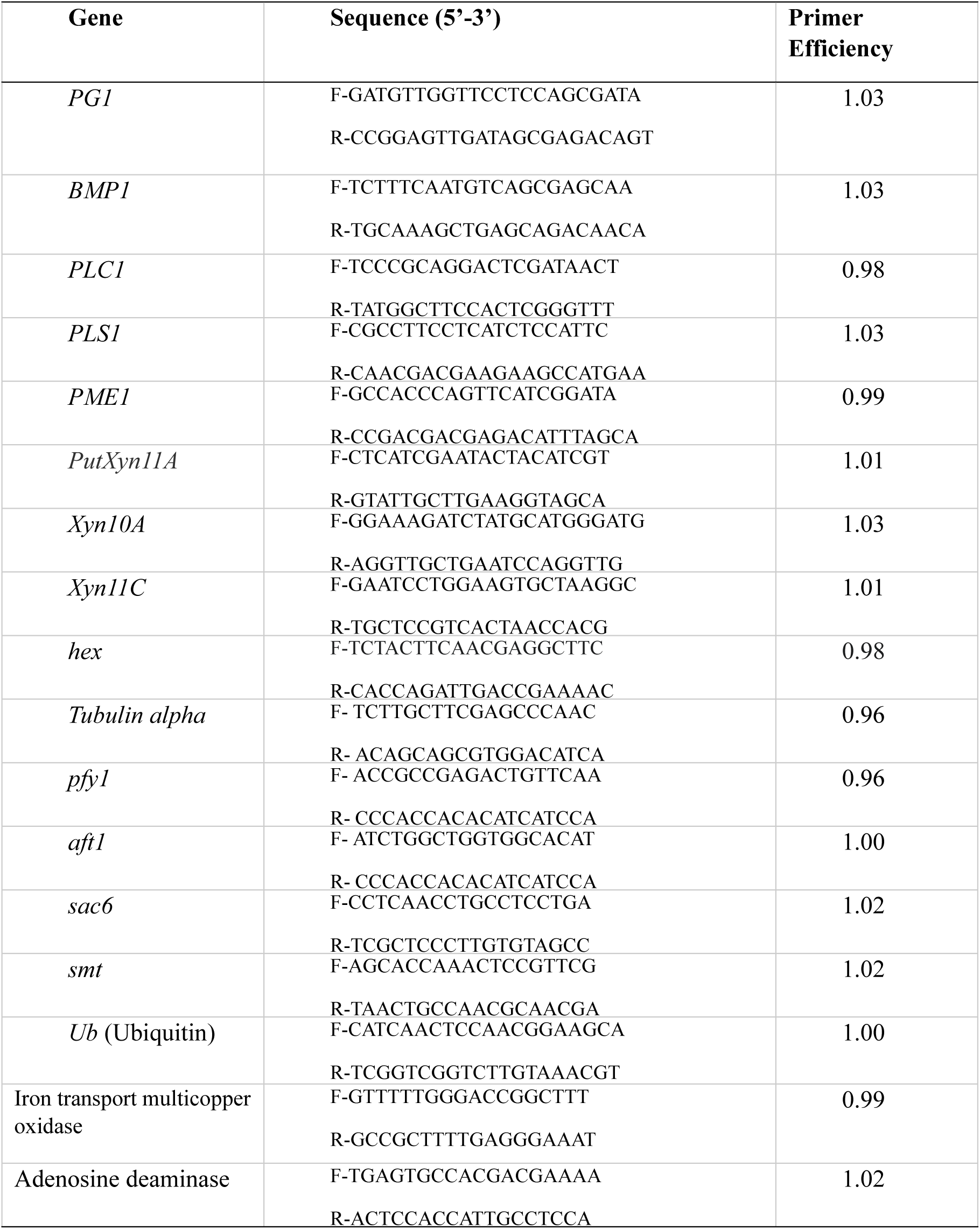
Primers used in RT-qPCR.

**Supplementary Table 3.**
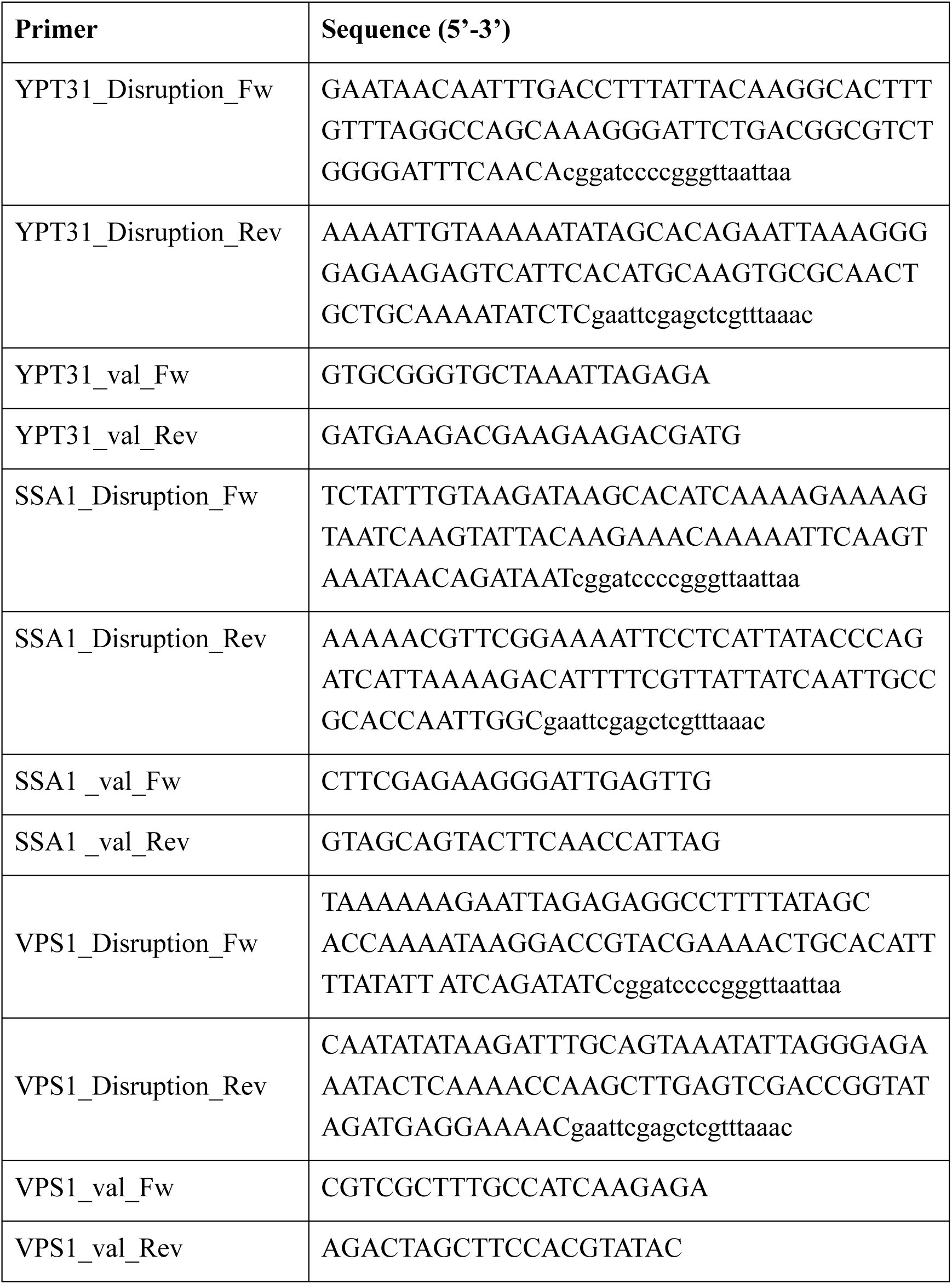
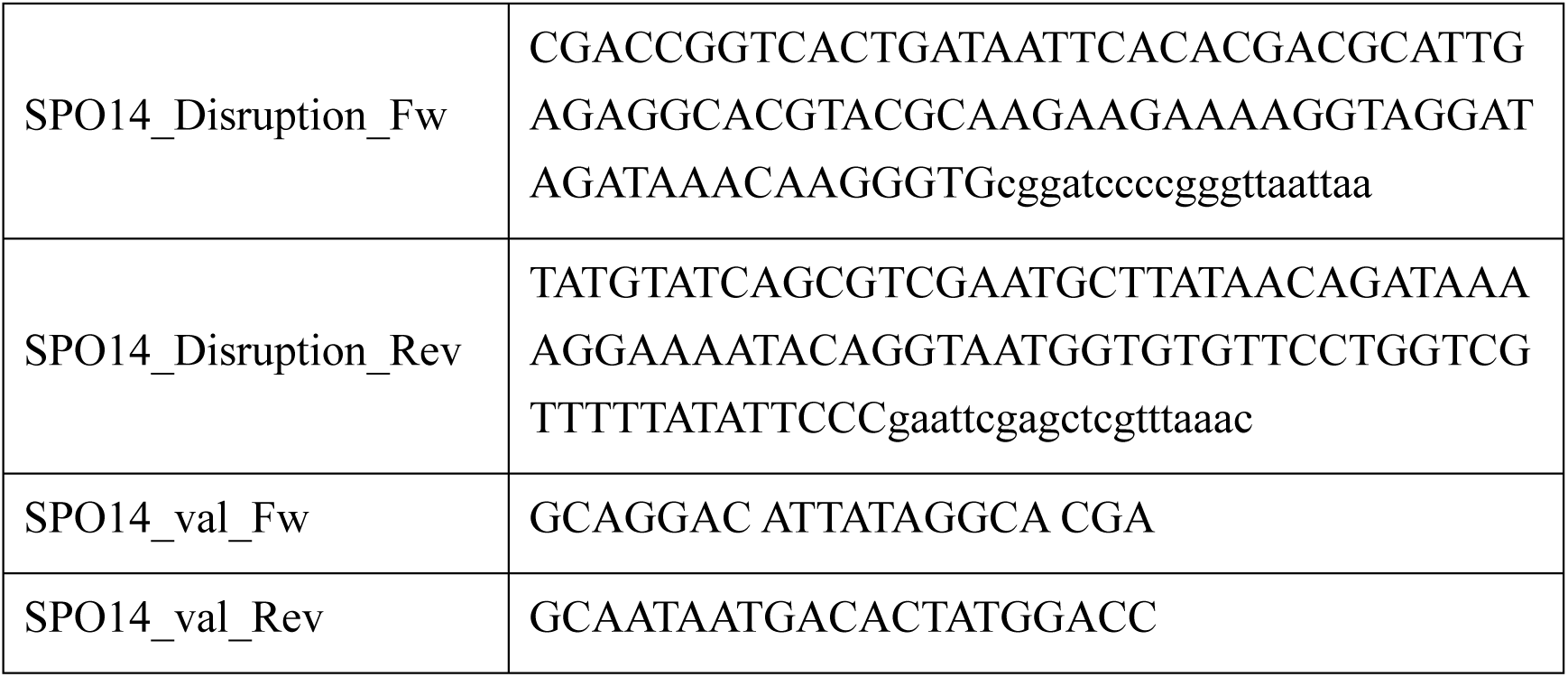
Oligonucleotides used for generating and validating *Saccharomyces cerevisiae* mutant strains.

**Supplementary Table 4.**
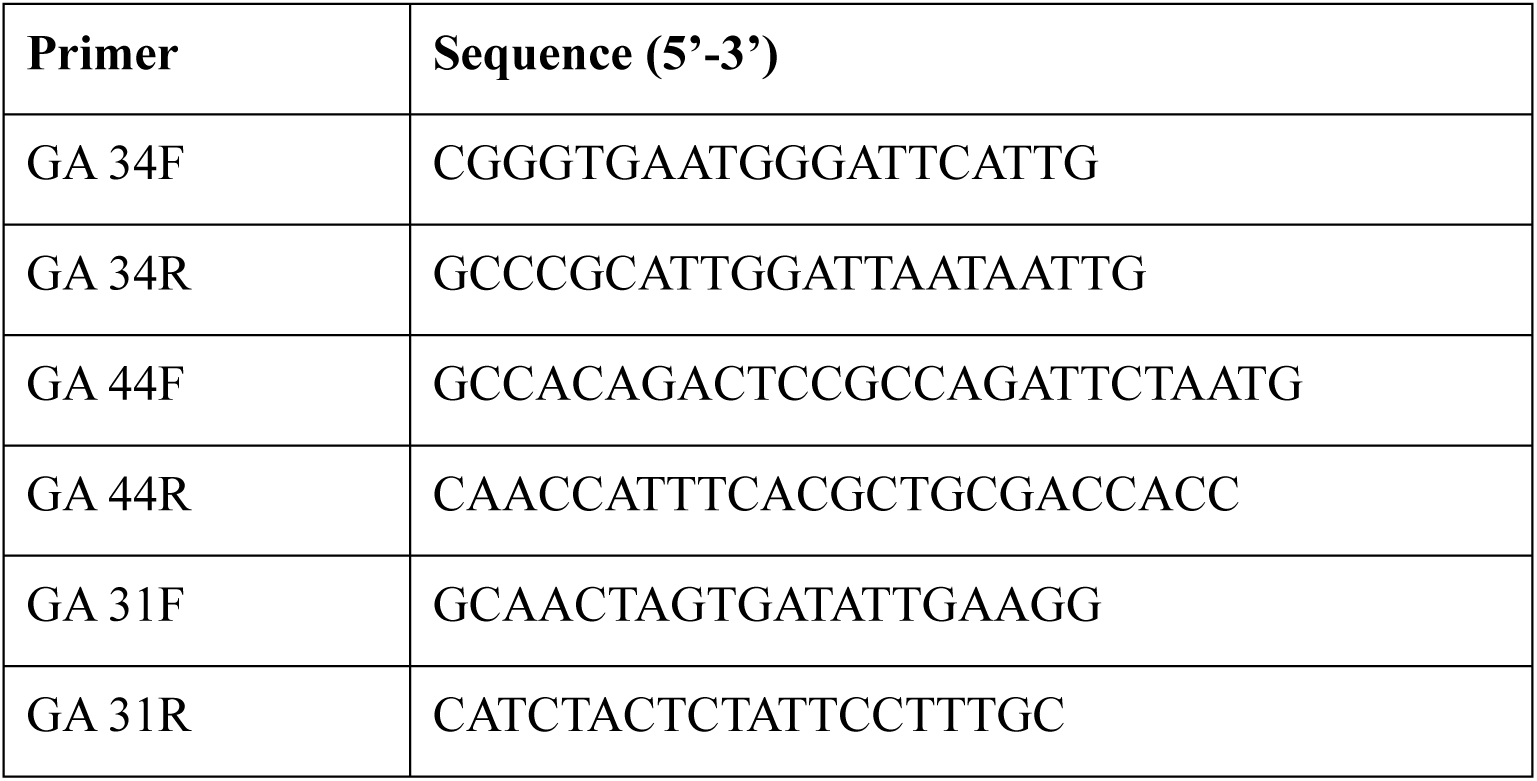
Oligonucleotides used for generating and validating *Botrytis cinerea* mutant strains.

## References

Aguayo C, Riquelme J, Valenzuela PDT, Hahn M, Silva Moreno E. 2011. Bchex virulence gene of Botrytis cinerea: characterization and functional analysis. Journal of General Plant Pathology 77: 230–238.

Argueso CT, Ferreira FJ, Epple P, To JPC, Hutchison CE, Schaller GE, Dangl JL, Kieber JJ. 2012. Two-component elements mediate interactions between cytokinin and salicylic acid in plant immunity. PLoS Genetics 8.

Arnold GRW. 2008. Farr, D. F.; Bills, G. F.; Chamuris, G. P.; Rossman, A. Y., Fungi on Plants and Plant Products in the United States. VIII, 1252 S. The American Phytopathological Society (APS) Press, St. Paul (Minnesota), 1989. ISBN 0-89054-099-3. Feddes Repertorium 101: 340–340.

Babosha A V. 2009. Regulation of resistance and susceptibility in wheat-powdery mildew pathosystem with exogenous cytokinins. Journal of Plant Physiology 166: 1892–1903.

Ballaré CL. 2011. Jasmonate-induced defenses: a tale of intelligence, collaborators and rascals. Trends in Plant Science 16: 249–257.

Bartnicki-Garcia S. 2002. Hyphal Tip Growth Outstanding Questions. *Published in* “Molecular Biology of Fungal Development” (H. D. Osiewacz, ed.).: 29–58.

Berepiki A, Lichius A, Read ND. 2011. Actin organization and dynamics in filamentous fungi. Nature Reviews Microbiology 9: 876–887.

Bolger AM, Lohse M, Usadel B. 2014. Trimmomatic: A flexible trimmer for Illumina sequence data. Bioinformatics 30: 2114–2120.

Boucrot E, Saffarian S, Massol R, Kirchhausen T, Ehrlich M. 2006. Role of lipids and actin in the formation of clathrin-coated pits. Experimental cell research 312: 4036–48.

Bradford MM. 1976. A rapid and sensitive method for the quantitation of microgram quantities of protein utilizing the principle of protein-dye binding. Analytical Biochemistry 72: 248–254.

Brito N, Espino JJ, González C. 2006. The endo-beta-1,4-xylanase xyn11A is required for virulence in Botrytis cinerea. Molecular plant-microbe interactions : MPMI 19: 25–32.

Buchfink B, Xie C, Huson DH. 2014. Fast and sensitive protein alignment using DIAMOND. Nature Methods 12: 59–60.

De Cal A, Garcia-Lepe R, Melgarejo P. 2000. Induced Resistance by Penicillium oxalicum Against Fusarium oxysporum f. sp. lycopersici: Histological Studies of Infected and Induced Tomato Stems. Am Phytopath Society.

Choi J, Choi D, Lee S, Ryu C-M, Hwang I. 2011. Cytokinins and plant immunity: old foes or new friends? Trends in Plant Science 16: 388–394.

Choi J, Huh SU, Kojima M, Sakakibara H, Paek K-HH, Hwang I. 2010. The cytokinin-activated transcription factor ARR2 promotes plant immunity via TGA3/NPR1-dependent salicylic acid signaling in arabidopsis. Developmental Cell 19: 284–295.

Conesa A, Götz S, García-Gómez JM, Terol J, Talón M, Robles M. 2005. Blast2GO: A universal tool for annotation, visualization and analysis in functional genomics research. Bioinformatics 21: 3674–3676.

Dean R, van Kan J, Pretorius Z, Hammond-Kosack K, Di pietro A, Spanu P, Redd J, Dickman M, Kahmann R, Ellis J, et al. 2012. The Top 10 fungal pathogens in molecular plant pathology. Molecular Plant Pathology 13: 414–430.

Dobin A, Davis CA, Schlesinger F, Drenkow J, Zaleski C, Jha S, Batut P, Chaisson M, Gingeras TR. 2013. STAR: Ultrafast universal RNA-seq aligner. Bioinformatics 29: 15–21.

Dub AM, Kokkelink L, Tudzynski B, Tudzynski P, Sharon A. 2013. Involvement of Botrytis cinerea small GTPases BcRAS1 and BcRAC in differentiation, virulence, and the cell cycle. Eukaryotic Cell 12: 1609–1618.

Ekena K, Vater CA, Raymond CK, Stevens TH. 1993. The VPS1 protein is a dynamin-like GTPase required for sorting proteins to the yeast vacuole. Ciba Foundation symposium 176.

Fillinger S, Elad Y. 2016. Botrytis -- the fungus, the pathogen and its management in agricultural systems.

Fischer-Parton S, Parton RM, Hickey PC, Dijksterhuis J, Atkinson HA, Read ND. 2000. Confocal microscopy of FM4-64 as a tool for analysing endocytosis and vesicle trafficking in living fungal hyphae. Journal of Microscopy 198: 246–259.

Frías M, González M, González C, Brito N. 2019. A 25-Residue Peptide From Botrytis cinerea Xylanase BcXyn11A Elicits Plant Defenses. Frontiers in Plant Science 10: 474.

Gentleman RC, Carey VJ, Bates DM, Bolstad B, Dettling M, Dudoit S, Ellis B, Gautier L, Ge Y, Gentry J, et al. 2004. Bioconductor: open software development for computational biology and bioinformatics. Genome biology 5.

Ghanem ME, Albacete A, Martínez-Andújar C, Acosta M, Romero-Aranda R, Dodd IC, Lutts S, Pérez-Alfocea F. 2008. Hormonal changes during salinity-induced leaf senescence in tomato (Solanum lycopersicum L.). Journal of Experimental Botany 59: 3039–3050.

Gomes CJ, Harman MW, Centuori SM, Wolgemuth CW, Martinez JD. 2018. Measuring DNA content in live cells by fluorescence microscopy. Cell Division 13: 6.

Gourgues M, Brunet-Simon A, Lebrun M-H, Levis C. 2004. The tetraspanin BcPls1 is required for appressorium-mediated penetration of Botrytis cinerea into host plant leaves. Molecular microbiology 51: 619–29.

Greene EM. 1980. Cytokinin production by microorganisms. The Botanical Review 46: 25–74.

Grosskinsky DK, Naseem M, Abdelmohsen UR, Plickert N, Engelke T, Griebel T, Zeier J, Novak O, Strnad M, Pfeifhofer H, et al. 2011. Cytokinins Mediate Resistance against Pseudomonas syringae in Tobacco through Increased Antimicrobial Phytoalexin Synthesis Independent of Salicylic Acid Signaling. Plant Physiology 157: 815–830.

Gupta R, Leibman-Markus M, Pizarro L, Bar M. 2020a. Cytokinin induces bacterial pathogen resistance in tomato. Plant Pathology In press.

Gupta R, Pizarro L, Leibman-Markus M, Marash I, Bar M. 2020b. Cytokinin response induces immunity and fungal pathogen resistance, and modulates trafficking of the PRR LeEIX2 in tomato. Molecular plant pathology: 10.1111/mpp.12978.

Hérivaux A, Dugé de Bernonville T, Roux C, Clastre M, Courdavault V, Gastebois A, Bouchara J-P, James TY, Latgé J-P, Martin F, et al. 2017. The Identification of Phytohormone Receptor Homologs in Early Diverging Fungi Suggests a Role for Plant Sensing in Land Colonization by Fungi. mBio 8.

Jameson PE. 2000. Cytokinins and auxins in plant-pathogen interactions – An overview. Plant Growth Regulation 32: 369–380.

Jedd G, Mulholland J, Segev N. 1997. Two new Ypt GTPases are required for exit from the yeast trans-Golgi compartment. The Journal of cell biology 137: 563–80.

Ji D, Chen T, Ma D, Liu J, Xu Y, Tian S. 2018. Inhibitory effects of methyl thujate on mycelial growth of Botrytis cinerea and possible mechanisms. Postharvest Biology and Technology 142: 46–54.

Jia Y, Li W. 2018. Phospholipase D antagonist 1-butanol inhibited the mobilization of triacylglycerol during seed germination in Arabidopsis. Plant diversity 40: 292–298.

Keshishian EA, Rashotte AM. 2015. Plant cytokinin signalling. Essays in biochemistry 58: 13–27.

Ketelaar T, Meijer HJG, Spiekerman M, Weide R, Govers F. 2012. Effects of latrunculin B on the actin cytoskeleton and hyphal growth in Phytophthora infestans. Fungal Genetics and Biology 49: 1014–1022.

Krantz KC, Puchalla J, Thapa R, Kobayashi C, Bisher M, Viehweg J, Carr CM, Rye HS. 2013. Clathrin coat disassembly by the yeast Hsc70/Ssa1p and auxilin/Swa2p proteins observed by single-particle burst analysis spectroscopy. The Journal of biological chemistry 288: 26721–30.

Laor D, Sade D, Shaham-Niv S, Zaguri D, Gartner M, Basavalingappa V, Raveh A, Pichinuk E, Engel H, Iwasaki K, et al. 2019. Fibril formation and therapeutic targeting of amyloid-like structures in a yeast model of adenine accumulation. Nature Communications 10: 1–11.

Lewis J, Papavizas G. 1987. Permeability changes in hyphae of Rhizoctonia solani induced by germling preparations of Trichoderma and Gliocladium. Physiology and Biochemistry 77: 699–703.

Li X, Gao M, Han X, Tao S, Zheng D, Cheng Y, Yu R, Han G, Schmidt M, Han L. 2012. Disruption of the phospholipase D gene attenuates the virulence of Aspergillus fumigatus. Infection and immunity 80: 429–40.

Liarzi O, Benichis M, Gamliel A, Ezra D. 2020. *trans* -2-Octenal, a single compound of a fungal origin, controls *Sclerotium rolfsii*, both *in vitro* and in soil. Pest Management Science 76: 2068–2071.

Liu Z, Bushnell WR. 1986. Effects of cytokinins on fungus development and host response in powdery mildew of barley. Physiological and Molecular Plant Pathology 29: 41–52.

Llanos A, François JM, Parrou JL. 2015. Tracking the best reference genes for RT-qPCR data normalization in filamentous fungi. BMC Genomics 16: 71.

Love MI, Huber W, Anders S. 2014. Moderated estimation of fold change and dispersion for RNA-seq data with DESeq2. Genome Biology 15.

Marhavý P, Bielach A, Abas L, Abuzeineh A, Duclercq J, Tanaka H, Pařezová M, Petrášek J, Friml J, Kleine-Vehn J, et al. 2011. Cytokinin Modulates Endocytic Trafficking of PIN1 Auxin Efflux Carrier to Control Plant Organogenesis. Developmental Cell 21: 796–804.

Mes J, Weststeijn E, Herlaar F. 1999. Biological and Molecular Characterization of Fusarium oxysporum f. sp. lycopersici Divides Race 1 Isolates into Separate Virulence Groups. Am Phytopath Society.

Mishina GN, Talieva MN, Babosha AV, Serezhkina GV, Andreev LN. 2002. Influence of phytohormones on development of conidial inoculum of causative agents of the phlox and barley powdery mildew | Vliianie fitogormonov na razvitie konidial’nogo inokuliuma vozbuditelei muchnistoi rosy floksa i iachmenia. Izvestiia Akademii nauk. Seriia biologicheskaia / Rossiiskaia akademiia nauk 29: 46–52.

Mok DW, Mok MC. 2001. Cytokinin metabolism and action. Ann. Rev. Plant Physiol. Plant Mol. Biol. 52: 89–118.

Momany M, Hamer JE. 1997. Relationship of actin, microtubules, and crosswall synthesis during septation in Aspergillus nidulans. Cell Motility and the Cytoskeleton 38: 373–384.

Motes CM, Pechter P, Yoo CM, Wang Y-S, Chapman KD, Blancaflor EB. 2005. Differential effects of two phospholipase D inhibitors, 1-butanol and N-acylethanolamine, on in vivo cytoskeletal organization and Arabidopsis seedling growth. Protoplasma 226: 109–123.

Naseem M, Philippi N, Hussain A, Wangorsch G, Ahmed N, Dandekar T, Dandekara T, Dandekar T. 2012. Integrated systems view on Networking by hormones in Arabidopsis immunity reveals multiple crosstalk for cytokinin. The Plant Cell 24: 1793–1814.

Nieto K, Frankenberger W. 1991. Microbial production of cytokinins. Soil biochemistry. Volume 6., 191-248. Bollag, J.-M. and G. Stotzky (eds.): Soil Biochemistry, Vol. 6 (Books in Soils, Plants, and the Environment Series). *Zeitschrift für Pflanzenernährung und Bodenkunde* 154: 74–74.

Peterson JR, Mitchison TJ. 2002. Small molecules, big impact: A history of chemical inhibitors and the cytoskeleton. Chemistry and Biology 9: 1275–1285.

Pfaffl MW. 2001. A new mathematical model for relative quantification in real-time RT-PCR. Nucleic Acids Res 29: e45.

Riquelme M, Aguirre J, Bartnicki-García S, Braus GH, Feldbrügge M, Fleig U, Hansberg W, Herrera-Estrella A, Kämper J, Kück U, et al. 2018. Fungal Morphogenesis, from the Polarized Growth of Hyphae to Complex Reproduction and Infection Structures. Microbiology and molecular biology reviews : MMBR 82.

Rudge SA, Zhou C, Engebrecht JA. 2002. Differential regulation of Saccharomyces cerevisiae phospholipase D in sporulation and Sec14-independent secretion. Genetics 160: 1353–1361.

Sakakibara H. 2006. Cytokinins: Activity, Biosynthesis, and Translocation. Annu Rev Plant Biol.

Schindelin J, Arganda-Carreras I, Frise E, Kaynig V, Longair M, Pietzsch T, Preibisch S, Rueden C, Saalfeld S, Schmid B, et al. 2012. Fiji: an open-source platform for biological-image analysis. Nature Methods 9: 676–682.

Schumacher J. 2012. Tools for Botrytis cinerea: New expression vectors make the gray mold fungus more accessible to cell biology approaches. Fungal Genetics and Biology 49: 483–497.

Schumacher J, Viaud M, Simon A, Tudzynski B. 2008. The Gα subunit BCG1, the phospholipase C (BcPLC1) and the calcineurin phosphatase co-ordinately regulate gene expression in the grey mould fungus Botrytis cinerea. Molecular Microbiology 67: 1027–1050.

Sharfman M, Bar M, Ehrlich M, Schuster S, Melech-Bonfil S, Ezer R, Sessa G, Avni A. 2011. Endosomal signaling of the tomato leucine-rich repeat receptor-like protein LeEix2. The Plant Journal 68: 413–423.

Sharma N, Rahman MH, Liang Y, Kav NN V. 2010. Cytokinin inhibits the growth of *Leptosphaeria maculans* and *Alternaria brassicae*. Canadian Journal of Plant Pathology 32: 306–314.

Silva-Moreno E, Brito-Echeverría J, López M, Ríos J, Balic I, Campos-Vargas R, Polanco R. 2016. Effect of cuticular waxes compounds from table grapes on growth, germination and gene expression in Botrytis cinerea. World Journal of Microbiology and Biotechnology 32: 74.

Sørensen JL, Benfield AH, Wollenberg RD, Westphal K, Wimmer R, Nielsen MR, Nielsen KF, Carere J, Covarelli L, Beccari G, et al. 2018. The cereal pathogen Fusarium pseudograminearum produces a new class of active cytokinins during infection. Molecular Plant Pathology 19: 1140–1154.

Staats M, van Kan JAL. 2012. Genome update of Botrytis cinerea strains B05.10 and T4. Eukaryotic Cell 11: 1413–1414.

Stravato VM, Buonaurio R, Cappelli C. 1999. First Report of *Fusarium oxysporum* f. sp. *lycopersici* Race 2 on Tomato in Italy. Plant Disease 83: 967–967.

Suzuki T, Miwa K, Ishikawa K, Yamada H, Aiba H, Mizuno T. 2001. The Arabidopsis sensor kinase, AHK4, can respond to cytokinin. Plant Cell Physiol. 42: 107–113.

Swartzberg D, Kirshner B, Rav-David D, Elad Y, Granot D. 2008. Botrytis cinerea induces senescence and is inhibited by autoregulated expression of the IPT gene. European Journal of Plant Pathology 120: 289–297.

Torres-Ossandón MJ, Vega-Gálvez A, Salas CE, Rubio J, Silva-Moreno E, Castillo L. 2019. Antifungal activity of proteolytic fraction (P1G10) from (Vasconcellea cundinamarcensis) latex inhibit cell growth and cell wall integrity in Botrytis cinerea. International Journal of Food Microbiology 289: 7–16.

Trapnell C, Roberts A, Goff L, Pertea G, Kim D, Kelley DR, Pimentel H, Salzberg SL, Rinn JL, Pachter L. 2012. Differential gene and transcript expression analysis of RNA-seq experiments with TopHat and Cufflinks. Nature Protocols 7: 562–578.

Tsahouridou PC, Thanassoulopoulos CC. 2002. Proliferation of Trichoderma koningii in the tomato rhizosphere and the suppression of damping-off by Sclerotium rolfsii. Soil Biology and Biochemistry 34: 767–776.

Valette-Collet O, Cimerman A, Reignault P, Levis C, Boccara M. 2003. Disruption of Botrytis cinerea pectin methylesterase gene Bcpme1 reduces virulence on several host plants. Molecular plant-microbe interactions : MPMI 16: 360–7.

Vásquez-Soto B, Manríquez N, Cruz-Amaya M, Zouhar J, Raikhel N V, Norambuena L. 2015. Sortin2 enhances endocytic trafficking towards the vacuole in Saccharomyces cerevisiae. Biological Research 48: 39.

Vater CA, Raymond CK, Ekena K, Howald-Stevenson I, Stevens TH. 1992. The VPS1 protein, a homolog of dynamin required for vacuolar protein sorting in Saccharomyces cerevisiae, is a GTPase with two functionally separable domains. Journal of Cell Biology 119: 773–786.

Vrabka J, Niehaus E-M, Münsterkötter M, Proctor RH, Brown DW, Novák O, Pěnčik A, Tarkowská D, Hromadová K, Hradilová M, et al. 2018. Production and Role of Hormones During Interaction of Fusarium Species With Maize (Zea mays L.) Seedlings. Frontiers in plant science 9: 1936.

Walker SK, Garrill A. Actin microfilaments in fungi.

Walters DR, McRoberts N. 2006. Plants and biotrophs: a pivotal role for cytokinins? Trends in plant science 11: 581–6.

Wang Y, Liu X, Chen T, Xu Y, Tian S. 2020. Antifungal effects of hinokitiol on development of Botrytis cinerea in vitro and in vivo. Postharvest Biology and Technology 159: 111038.

Wellman F. 1939. A technique for sutdying host resistance and pathogenicity in tomato Fusarium wilt. Phytopathology.

Werner T, Schmülling T. 2009. Cytokinin action in plant development. Current opinion in plant biology 12: 527–38.

Williamson B, Tudzynski B, Tudzynski P, Van Kan JAL. 2007. Botrytis cinerea: The cause of grey mould disease. Molecular Plant Pathology 8: 561–580.

Xie C, Mao X, Huang J, Ding Y, Wu J, Dong S, Kong L, Gao G, Li CY, Wei L. 2011. KOBAS 2.0: A web server for annotation and identification of enriched pathways and diseases. Nucleic Acids Research 39.

Zheng L, Campbell M, Murphy J, Lam S, Xu JR. 2000. The BMP1 gene is essential for pathogenicity in the gray mold fungus Botrytis cinerea. Molecular plant-microbe interactions : MPMI 13: 724–32.

Žižková E, Dobrev PI, Muhovski Y, Hošek P, Hoyerová K, Haisel D, Procházková D, Lutts S, Motyka V, Hichri I. 2015. Tomato (Solanum lycopersicum L.) SlIPT3 and SlIPT4 isopentenyltransferases mediate salt stress response in tomato. BMC Plant Biology 15: 1–20.

